# Clathrin adaptors drive phase separation in endocytosis and trafficking

**DOI:** 10.1101/2025.11.20.689461

**Authors:** George Draper-Barr, Janina Schiller, Katharina Veith, Stephan Niebling, David Ruiz-Carrillo, Ziqiang Huang, Christian Tischer, Roland Thuenauer, Lucas A. Defelipe, Maria García-Alai

**Author notes:** Corresponding authors: Maria García-Alai and Lucas A. Defelipe. equally contributed (George Draper-Barr, Janina Schiller).

## Abstract

Liquid–liquid phase separation (LLPS) underlies the formation of biomolecular condensates that organize cellular processes, including endocytosis and membrane trafficking. Although LLPS has been implicated in clathrin-mediated pathways, the specific contribution of adaptor proteins to condensate formation and function remains unclear. Here we show that two yeast adaptors, Ent5 and Sla2, undergo LLPS in vitro and that this property correlates with their distinct roles in membrane trafficking and condensate recruitment in vivo. Ent5 condensation is driven by a dynamic helix within its disordered region that acts as a molecular “sticker.” Deletion of this helix disrupts membrane-associated condensation and leads to altered cargo trafficking dynamics in vivo, with Ent5-dependent transport events becoming delayed. In contrast, Sla2 behaves both as a driver and as a client of LLPS, with its coiled-coil (CC) region mediating condensation. Together, these findings reveal that endocytic adaptors can promote condensate formation through distinct structural features, thereby coupling clathrin binding and membrane association through phase separation.

## Introduction

Biological condensates form in cells when macromolecules, primarily proteins or nucleic acids, undergo liquid-liquid phase separation (LLPS), transitioning from an initially solubilized form into at least two distinct phases. This process is driven by a shift in interaction preferences: from macromolecule-water interactions to water-water and macromolecule-macromolecule interactions. *In vitro*, the simplest case of LLPS involves the formation of a dense phase in the form of round droplets, surrounded by a dilute aqueous environment(1). These dense droplets retain liquid-like properties, allowing for molecular exchange with the dilute phase, fusion with other droplets, and growth(2). Although LLPS is described as a spontaneous phenomenon, it is tightly regulated by the unique characteristics of each macromolecule, as well as by solvent composition, macromolecular concentration, temperature, and other environmental factors(1, 3, 4).

When LLPS-formed, protein-rich condensates come into close contact with the surface of the cell or organelle membrane, they can adhere to it, “wetting” the membrane and leading to its deformation. This interaction plays a key role in biologically relevant processes such as budding, nanotube formation, compartmentalization, and membrane fission(5). Moreover, LLPS driven by intrinsically disordered proteins (IDPs) can exert steric pressure, due to their large hydrodynamic radii, sufficient to induce membrane bending(6).

IDPs play diverse functional roles in biomolecular condensates. They can drive phase separation, tune the physical and biochemical properties of condensates, suppress aberrant aggregation, and act as regulatory elements that modulate condensate behavior in response to post-translational modifications(6). IDPs contain “*stickers and spacers”: stickers* are responsible for percolation and form reversible interactions through specific residues or motifs in intrinsically disordered regions (IDRs). These may include inducible secondary structural elements, such as helices or beta strands, which can further influence percolation and network stability(7). *Spacers*, in contrast, modulate solubility and influence sticker-sticker cooperativity. IDRs contribute uniquely to LLPS by enabling multivalent molecular interactions with a high degree of regulatory flexibility(8, 9). In addition, Ramirez et al(10) have shown that coiled-coil (CC) structures possess physical features that enable them to drive the LLPS of proteins. CC-containing proteins can potentially utilize both polymeric and multimeric multivalency mechanisms to promote LLPS through their CC domains. Among these, multimerization has a strong effect on LLPS propensity, while polymer multivalency has a weaker influence.

When LLPS is coupled with percolation, specific interactions facilitate the formation of small percolating clusters composed of one or more types of macromolecules(9, 11). Macromolecules essential for LLPS are termed *drivers*, as they can undergo phase separation independently. In contrast, *clients* are molecules that do not phase-separate on their own but can partition into condensates formed by drivers(12, 13).

Clathrin-mediated endocytosis requires the concerted action of membrane-binding proteins, known as adaptors, which link the membrane, the clathrin scaffold, and the actin polymerization machinery. This coordinated interaction results in membrane invagination and vesicle formation. Ede1, the yeast homolog of mammalian Eps15, is an early-arriving endocytic protein and a key initiation factor, although it does not bind directly to clathrin. In the absence of Ede1, most other early endocytic proteins lose their punctate localization, and endocytic uptake is significantly reduced. When present above a threshold concentration, Ede1 forms condensates that recruit other endocytic proteins and exhibit properties characteristic of phase-separated liquid droplets. The central region of Ede1, which contains a CC domain, is essential for both condensate formation and Ede1’s function in endocytosis(14).

In order to understand whether the clathrin scaffold could function as a lattice embedded within a phase-separating environment, and which adaptors might nucleate or regulate this process, we conducted a screening experiment to investigate the role of LLPS in clathrin-mediated endocytosis and trafficking proteins. We selected several clathrin-binding adaptors based on their N-terminal domain interactions with clathrin heavy chain (NTD-CHC), as defined by Defelipe et al.(15), and with clathrin light chain (CLC)(16).

In this study, we report that Ent5, an epsin involved in trafficking, and Sla2, a key component of the endocytic mid-coat, both undergo LLPS. We further characterize their roles as drivers and clients. Ent5 consists of an Epsin N-Terminal Homology (ENTH) domain, which binds to phosphatidylinositol-3,5-bisphosphate [PtdIns(3,5)P₂](17, 18), followed by an IDR(19). We demonstrate that Ent5 promotes phase separation through a *sticker*-inducible α-helix and is capable of recruiting clathrin and Ent3. In contrast, Sla2, the yeast homolog of mammalian Hip1r, can be recruited by Ede1 condensates(14) as a client but also induces LLPS (behaves as a driver). For this second role, phase separation would be mainly mediated by its CC region, particularly in the context of a membrane environment.

## Results

### Ent5 and Sla2 form liquid condensates *in vitro*

The clathrin triskelion is a three-legged structure composed of three clathrin heavy chains (CHC) and three clathrin light chains (CLC). It self-assembles into a polyhedral lattice, forming a scaffold on the inner surface of the cell membrane: the clathrin coat(20). The N-terminal domain of CHC (NTD-CHC) is responsible for interacting with proteins that contain a Clathrin Binding Motif (CBM)(15). CHC lacks significant intrinsically IDRs and, accordingly, is not predicted to form condensates spontaneously. In contrast, CLC contains an unstructured region known to interact with Sla2(16) and a long alpha-helix that binds to the distal domain of CHC. Unlike CHC, CLC is predicted to spontaneously undergo phase separation (see Table 1).

**Table 1.** Proteins involved in endocytosis and trafficking. ^1^Prediction from FuzDrop/ Probability of Spontaneous LLPS > 0.6 is considered to form condensates.

| Name | UniProtID | IDR | Membrane Binding | Function | LLPS Prediction<sup>1</sup> |
| --- | --- | --- | --- | --- | --- |
| Apl2 | P36000 | Yes | Yes | AP1 Complex - TGN Trafficking | No |
| Apl5 | Q08951 | Yes | Yes | AP3 Complex - Golgi | No |
| CHC | P22137 | No | No | Formation of Protein Coat | No |
| CLC | P17891 | Yes | No | Regulation of Coat Assembly | Yes |
| Ede1 | P34216 | Yes | No | Endocytosis initiation factor | Yes |
| Ent1 | Q12518 | Yes | Yes | Adaptor protein in Endocytosis | Yes |
| Ent2 | Q05785 | Yes | Yes | Adaptor protein in Endocytosis | Yes |
| Ent3 | P47160 | Yes | Yes | Adaptor protein in TGN Trafficking | Yes |
| Ent5 | Q03769 | Yes | Yes | Adaptor protein in TGN Trafficking | Yes |
| Gga1 | Q06336 | Yes | No | TGN Trafficking | No |
| Gga2 | P38817 | Yes | No | TGN Trafficking | Yes |
| Sla1 | P32790 | Yes | No | Cytoskeletal protein binding protein; required for assembly of the cortical actin cytoskeleton | Yes |
| Sla2 | P33338 | Yes | Yes | Connect the membrane with the actin cytoskeleton | No |
| Swa2 | Q06677 | Yes | No | Cofactor for the uncoating of clathrin-coated vesicles (CCVs) by Hsp70-type chaperones | Yes |
| Yap1801 | P38856 | Yes | Yes | Clathrin Adaptor Protein | Yes |
| Yap1802 | P53309 | Yes | Yes | Clathrin Adaptor Protein | Yes |

To test the hypothesis that clathrin adaptors are prone to induce LLPS, we performed a combined analysis using AI-based modeling for IDR prediction and *in vitro* phase separation screening of adaptors known to bind clathrin and participate in endocytosis or trans-Golgi network (TGN) trafficking. Our screening identified several proteins with the potential to undergo phase separation, based on predictions from FuzDrop(21, 22), with a probability of spontaneous LLPS greater than 0.6 (see Table 1 and Supplementary Table S1). Supplementary Table S1 includes information about the IDRs, specifically highlighting the “Droplet-promoting regions”, those predicted residues to be involved in intramolecular interactions such as percolation. These regions could potentially act as LLPS “*stickers*.”

Ede1, known to form condensates spontaneously due to its extensive IDRs(14), was detected as a positive hit in our screening. The adaptor proteins Ent1, Ent2, YAP1801, and YAP1802 share structurally related N-terminal membrane-binding domains(23) and C-terminal IDRs containing CBMs(15). All four proteins are predicted to spontaneously form condensates (see Table 1).

Sla2 contains an ANTH domain at its N-terminal region, followed by an IDR, a CC-region, a force-sensing domain (REND), an actin-binding domain (THATCH), and a regulatory helix (see Supplementary Fig S1). Despite possessing an IDR, Sla2 is not predicted to spontaneously phase separate. In contrast, Sla1, which interacts with Sla2 to regulate actin polymerization, contains three SH3 domains, a membrane-interacting PH domain, and a large IDR(16) is predicted to spontaneously phase separate (Table 1).

Among other targets predicted to undergo spontaneous phase separation, we identified Swa2, an auxilin-like protein involved in clathrin uncoating. Swa2 includes three TPR domains (which interact with Hsp70-type chaperones), a J domain that stimulates the ATPase activity of the chaperone, and several IDRs.

Regarding proteins involved in trafficking, Ent3 and Ent5 both contain ENTH domains at their N-termini and IDRs at their C-terminal regions, similar to Ent1 and Ent2. However, Ent3 and Ent5 differ functionally, being involved in TGN trafficking and recognizing PI(3,5)P₂ rather than PI(4,5)P₂. Both Ent3 and Ent5 are predicted to spontaneously phase separate.

Gga1 and Gga2, trans-Golgi adaptor proteins, recognize linear motifs (DxxLL) through their Vps27, Hrs, and STAM (VPS) domains and contain Gamma-adaptin Ear (GAE) domains that mediate protein-protein interactions. Although both possess long IDRs, Gga2 is predicted to spontaneously phase separate, while Gga1 is not. This difference can be partly explained by droplet-promoting predictions from FuzDrop: whereas Gga2 has long droplet-promoting regions (Supplementary Table S1), Gga1 lacks them.

Lastly, Apl2 (AP-1 subunit beta) and Apl5 (AP-3 subunit delta) both contain multiple Armadillo-like repeats and relatively short IDRs. Neither Apl2 nor Apl5 is predicted to spontaneously phase separate, likely due to the insufficient length and biophysical properties of their IDRs.

We then conducted a chemical screening by varying biophysical parameters such as temperature, ionic strength, pH, and the presence of crowding agents focused on the top-scoring targets from Table 1 and those for which we could identify structural regions likely to act as stickers (see Supplementary Table S1). Our experiments identified Ent5 and Sla2 as capable of inducing LLPS *in vitro* (see Fig 1).

**Figure 1.**
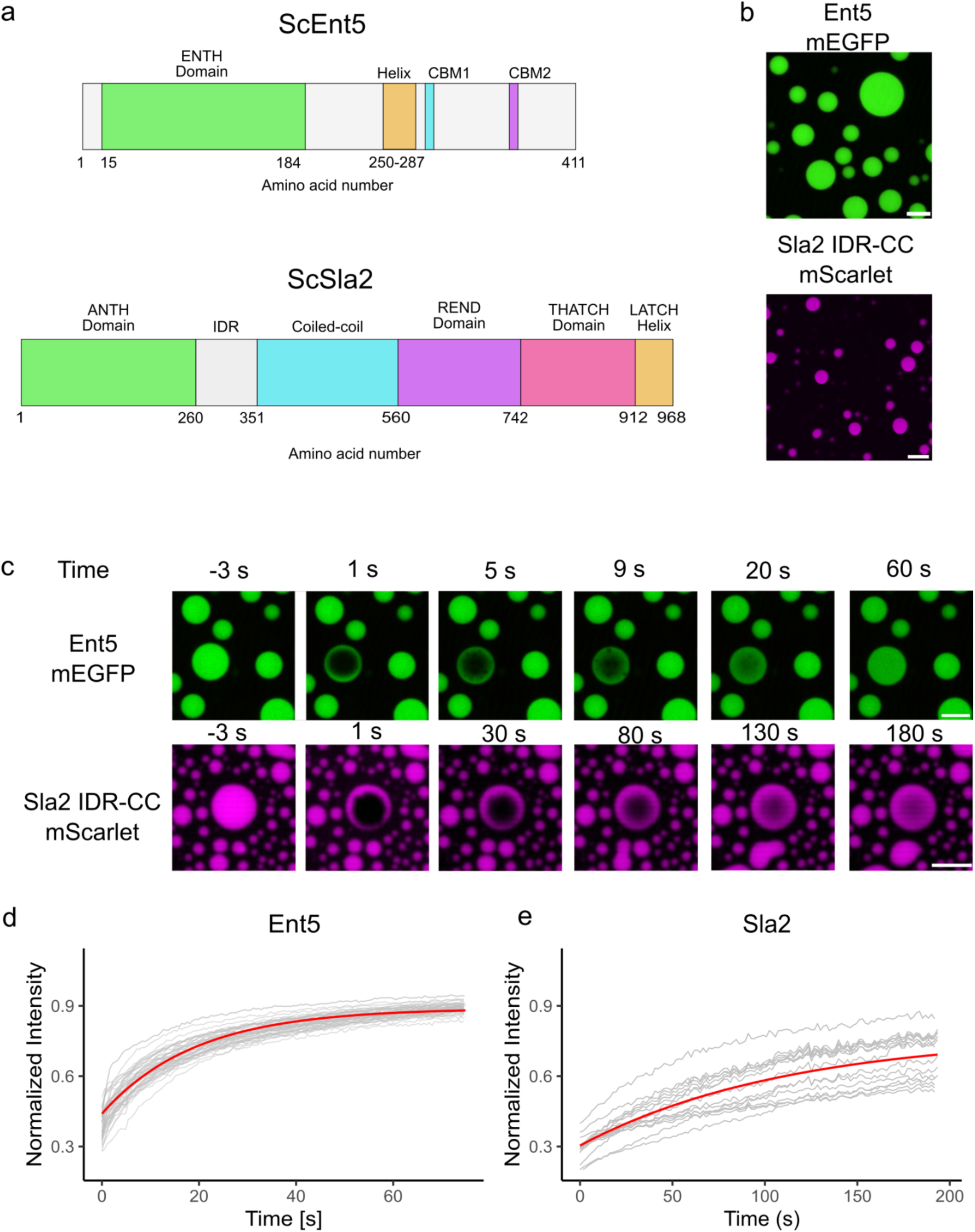
Ent5 and Sla2 form liquid condensates. a) Domain architecture of Ent5 and Sla2 from *Saccharomyces cerevisiae*. Ent5 is formed by a membrane binding ENTH domain (15-184) and a IDR (211-411), which is composed of a predicted central helix (249-287), Clathrin Binding Motif (CBM) 1 (LIDL, 293-296) and 2 (LLDW, 355-358). Sla2 is composed of a membrane binding ANTH domain (1-260), IDR (260-351), Coiled-coil (CC,351-560), REND domain (560-742), THATCH domain (742-912) and a LATCH helix (912-968). b) Exemplar images of condensates at room temperature (21 °C) for Ent5 full-length N-terminal tagged with mEGFP (upper panel) and a truncated Sla2 (IDR and CC) tagged C-terminally with mScarlet (bottom panel). Ent5 droplets formed at 18 µM protein concentration in buffer (100 mM Tris pH 8, 20 mM NaCl, and 0.5 mM TCEP). Sla2 droplets formed at 25 µM protein concentration in the presence of PEG (2,5 % PEG 8000, 30 mM HEPES pH 8, 150 mM NaCl, 0.5 mM TCEP). The white scale bar represents 5 µm. c) Representative FRAP time course of droplets for Ent5-mEGFP (upper panels) and Sla2 IDR-CC-mScarlet (lower panels). Upper panels show confocal images of an Ent5-mEGFP droplet before photobleaching (–3 s) and at 1, 5, 9, 20, and 60 s after bleaching; lower panels show confocal images of a Sla2 IDR-CC-mScarlet droplet before photobleaching (-3 s) and at 1, 30, 80, 130, and 180 s after bleaching. d-e) Normalized intensity recovery plots for individual droplets are shown in grey, n=45 for Ent5 (panel d) and n=24 for Sla2 (panel e). A simple diffusion model fit is overlaid in red. Ent5 recovered with a time constant of (19.2 ±0.2) s (d). Sla2 recovered with a time constant of (112.2 ± 2.4) s (e).

The AlphaFold3 model of Ent5 contains the predicted ENTH domain and a long IDR (see Fig 1A and 2A). Interestingly, a well-predicted helix with an average predicted Local Distance Difference Test (pLDDT) of 74.36 (see Fig 2B) is observed within the IDR and is not present in Ent3 (see Supplementary Fig S2). Ent5 is found only in Phyla Ascomycota and Basidiomycota Dikarya (higher fungi, including *S. cerevisiae*) and has been implicated in the trafficking of chitin synthase Chs3 (UniProt ID P29465) (24). The helix is located upstream of the clathrin-binding motifs CBM1 and CBM2(24) (see Fig 1a and 2b).

**Figure 2.**
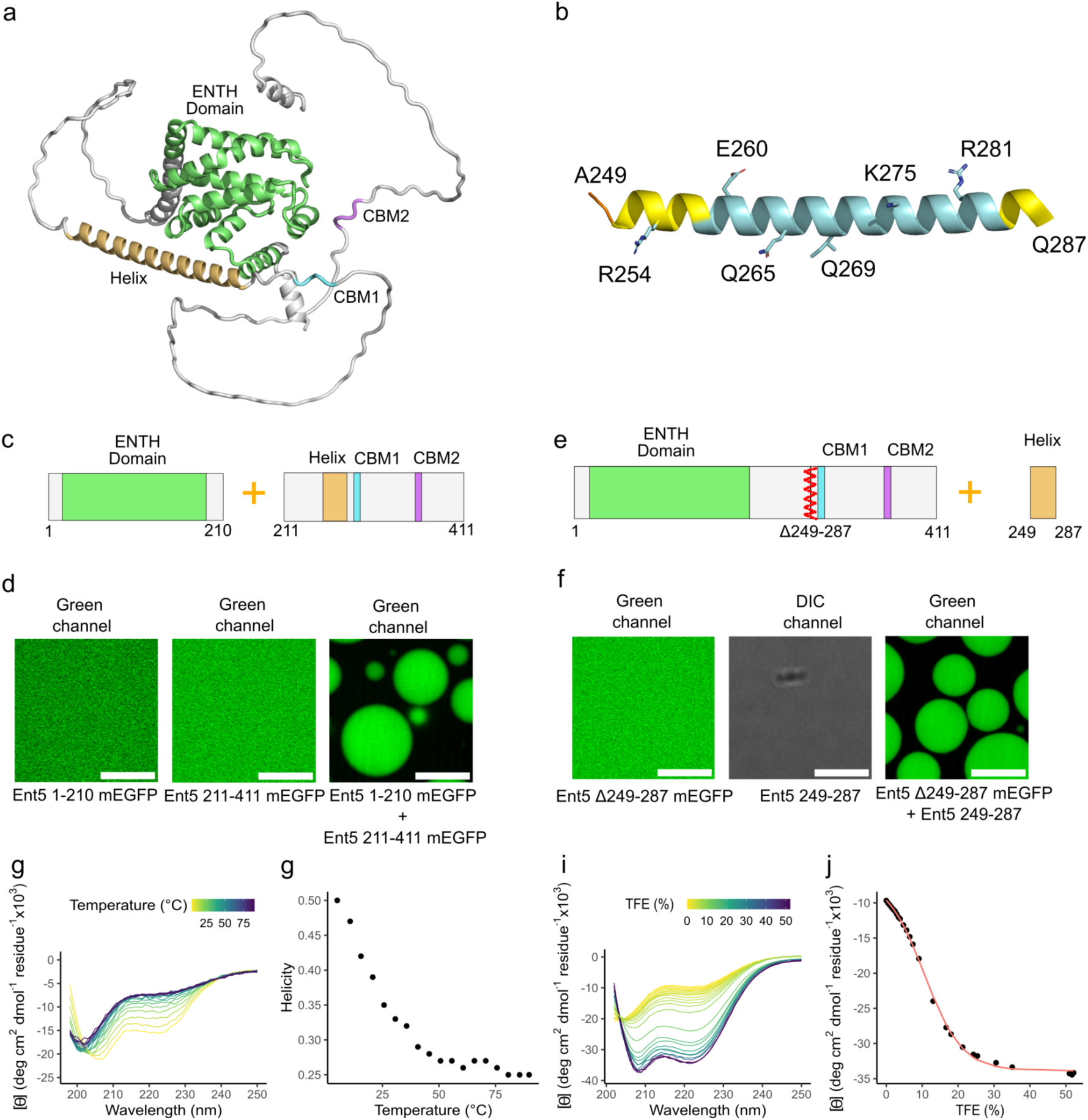
Ent5 ENTH Domain and IDR Helix Complement Each Other to Form Condensates. a) AlphaFold model of Ent5: In green the ENTH domain, the predicted helix in orange, CBM1 in cyan and CBM2 in purple. b) Zoom into the predicted helix. In light blue the model confidence prediction is high (90 > pLDDT > 70) and lower in yellow (70 > pLDDT > 50). c) Constructs used for the complementation experiment: ENTH domain (residues 1-210) and the IDR region (residues 211-411). d) Ent5 ENTH domain (16 µM, left panel) and the IDR region (16 µM, middle panel) in 100 mM Tris (pH 8), 20 mM NaCl, and 0.5 mM TCEP at room temperature (21 °C) do not phase separate. When the two constructs are mixed (both at 16 µM) the sample forms condensates (right panel). e) Constructs used for the complementation experiment: Ent5 ΔHelix (mutant with a deletion of residues 249-287) and a peptide for residues 249-287 (helix). f) Ent5 ΔHelix (16 µM, left panel) and helix (520 µM, right panel) do not phase separate (same buffer conditions as used in d). When the two constructs are mixed at these concentrations the sample forms condensates (right panel). g) Circular Dichroism (CD) spectra of the Ent5 Helix peptide (helix) were recorded over a temperature ramp from 5-90 °C. h). The percentage contribution of helical secondary structure,calculated from the molar ellipticity spectra using the Peptide Helicity module of ChiraKit(41),is plotted against temperature. i) Far UV CD spectra for the helix were recorded in the presence of TFE titrated from 0 to 52% (v/v) in 10 mM Tris (pH 7.5). j) Molar ellipticity at 222 nm plotted against TFE percentage. Data were fitted using the Custom Analysis module of ChiraKit(41) with Eq. 2 the following parameters (and their corresponding percentual errors calculated as absolute values): θTFE = -6470.5 ± 532.7, θwater = -33867 ± 159.5, ΔGwater = -1.19 ± 0.07 kcal/mol, m = 11.23 ± 0.44. White scale bars (5 µm) in panels b and d indicate the scale for all microscopy images.

Ent5 undergoes phase separation in response to changes in salt concentration and the presence of crowding agents (Fig 1) at low micromolar concentrations. To confirm that the phase separation behavior of Ent5 is not due to the fluorescent tag, we performed Differential interference contrast (DIC) imaging without the mEGFP fusion (see supplementary Fig 3). During visualization, frequent fusion events between individual droplets were observed, indicating their liquid nature. To assess the liquidity of the condensates, Fluorescence Recovery After Photobleaching (FRAP) was performed. Briefly, individual droplets within the field of view were selectively photobleached using a high-intensity laser, and fluorescence intensity was recorded over time to monitor recovery. The rate and uniformity of fluorescence return reflect the mobility of fluorescent molecules, consistent with a liquid-like state (Fig 1b and c). The recovery kinetics were fitted using a simple diffusion model, with a fast recovery of fluorescence (Tau: ∼19 s).

To identify the protein regions involved in LLPS, we designed two constructs of Ent5: one corresponding to the ENTH domain and one to the IDR containing the helix (Fig 2c). Our results indicate that only the full-length protein undergoes LLPS. Interestingly, mixing the ENTH domain and IDR constructs (at concentrations at which the full-length protein phase separates) results in LLPS, suggesting that the two domains complement each other (Fig 2c). However, the individual constructs do not form droplets under the same conditions.

In order to test the existence of the AlphaFold-predicted helix experimentally, we performed Far-UV Circular Dichroism (CD) analysis on a peptide containing residues 249-287. Our results show that the peptide is primarily alpha-helical in solution (Fig 2g-j). A temperature ramp experiment revealed a decrease in helical content, with a maximum of 50% helix observed at 5 °C. In the presence of Trifluoroethanol (TFE), the helix was further stabilized, reaching 98% helical content. Proline substitutions in the peptide (see Methods), which disrupt alpha-helices, significantly reduced the helical propensity of the region (see Supplementary Fig 4).

To test whether the helix would behave as a *sticker* and therefore be responsible for LLPS, we generated a deletion mutant (Ent5 ΔHelix) (Fig 2e). Only when Ent5 ΔHelix is supplemented with higher concentrations of the helix peptide 249-287 is LLPS restored, indicating that the helix functions as a *sticker* promoting phase separation. (see Fig 2f).

It has been shown that Sla2 is recruited to phase-separated Ede1 condensates in *S. cerevisiae*, and this has been proposed to occur via CC-interactions(14). Here, we demonstrate that a full-length Sla2 construct forms condensates from ∼2.5 µM in the presence of crowding agents (Fig 3a).

**Figure 3.**
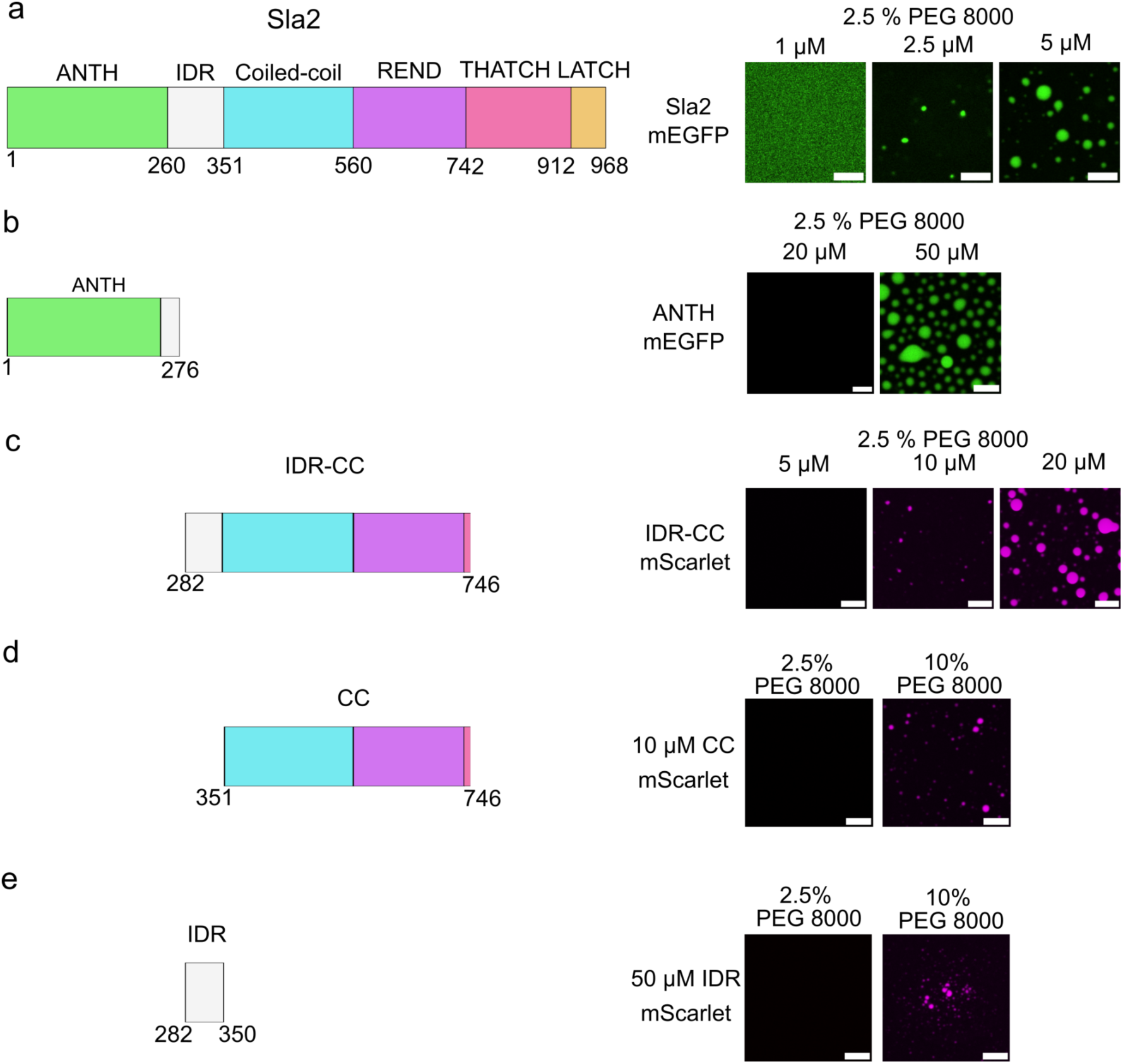
Condensate formation of Sla2 can be triggered by different regions. a) Sla2-mEGFP forms condensates at 2.5 μM (smaller droplets) and at 5 μM (larger droplets). Experiments were performed at room temperature (RT, 21 °C) in 30 mM HEPES pH 8, 150 mM NaCl, and 0.5 mM TCEP. b) Sla2 ANTH-mEGFP domain forms droplets at 50 μM in the presence of 2.5% (w/v) PEG 8000, in 30 mM HEPES pH 8, 150 mM NaCl, 0.5 mM TCEP at RT. c) Sla2 IDR-CC–mScarlet (residues 282-746) forms condensates at 10 μM (smaller droplets) and at 20 μM (larger droplets) under the same PEG buffer conditions as in b. d) Sla2 CC-mScarlet (residues 351-746) forms droplets in the presence of 10% (w/v) PEG 8000 at RT. (e) Sla2 IDR-mScarlet (residues 282-350) forms condensates at 50 μM in the presence of 10% (w/v) PEG 8000. White scale bars in panels a-e represent 5 µm.

To identify the protein regions responsible for LLPS, we designed truncated Sla2 constructs and tested them *in vitro* (see Fig 3). The CC region also includes the REND domain, previously implicated in forming independent clusters(25). Each region was expressed with a C-terminal fluorescent tag (either mEGFP or mScarlet). The Sla2 ANTH domain forms condensates at 20-50 µM, while the Sla2 IDR-CC construct forms larger, fluid condensates at a lower threshold (∼10 µM). The IDR and CC constructs form condensates only in 10% (W/V) PEG 8000, at concentrations of 50 µM and 10 µM, respectively. Our results indicate that the central region of Sla2 (IDR-CC), composed of the IDR (residues 286-350) and the CC region (residues 351-746), forms liquid condensates in the presence of 2.5% PEG 8000. These condensates exhibit uniform fluorescence recovery in FRAP experiments, confirming their fluid nature (Fig 1c). Similar to the complementation observed between the ENTH domain and IDR in Ent5 (Fig 2), addition of the Sla2 ANTH domain to the IDR-CC construct results in a synergistic enhancement of Sla2 condensation (Supplementary Fig 5). The droplet-to-solution fluorescence intensity ratio (ID/IS) for Sla2 IDR-CC in the presence of the membrane-binding domain is 15 at 5 µM [ANTH] and 10 at 20 µM [ANTH]. These values are several-fold higher than those observed for Sla2 IDR-CC alone. Additionally, ANTH domain ID/IS increases from 5.5 to 7.4. In conclusion, the combination of the ANTH and IDR-CC domains exhibits an enhanced capacity for partitioning into condensates.

### Ent5 behaves as an LLPS *driver*

To investigate the biophysical properties of Ent5 in LLPS, we analyzed its kinetics of condensation *in vitro*. Mass photometry was used to monitor the presence of nanoclusters in equilibrium at low (5 mM) and high (100 mM) NaCl concentrations (see Supplementary Fig S6). Ent5 monomeric peaks (∼91-97 kDa) were observed under all conditions. However, the wild-type construct exhibited higher molecular weight (MW) species (>200 kDa), corresponding to nanoclusters that appeared exclusively at low salt concentrations. In contrast, Ent5 ΔHelix showed significantly less signal above 200 kDa.

To further investigate the transition from monomers to higher oligomeric forms and to understand the kinetics of this assembly process, we performed Stopped-Flow light scattering and Stopped-Flow Small-Angle X-ray Scattering (SAXS) experiments (see Supplementary Figure S7). We aimed to quantify the speed of this transition and identify potential growth phases during nanocluster formation. We initiated the assembly by rapidly mixing Ent5, initially stabilized in a high-salt buffer, with a low-salt buffer to trigger the formation of larger molecular weight (MW) species. Both light scattering and SAXS are suitable to probe nanocuster formation - covering different size regimes of nanoclusters. Light absorption measurements at 600 nm (Supplementary Fig 7) revealed a biexponential kinetic profile, characterized by a rapid initial phase (τ₁ = 108 ± 7 ms) followed by a slower secondary phase (τ₂ = 843 ± 42 ms). This suggests that the assembly process proceeds through at least two distinct kinetic steps. To further characterize the emergence of nanoclusters, we employed SAXS experiments. The SAXS trace in the Guinier region displayed complex behavior, which could not be fitted with a simple exponential curve. However, a qualitative analysis indicated a half-time of approximately 200 ms, suggesting a rapid and structured growth phase.

Ent5 is a main player in yeast endosomal trafficking and contains two predicted Clathrin binding motifs (see Fig 1a). One of the binding sites, CBM1, has been shown to form a low micromolar complex with the N-terminal domain of Clathrin Heavy Chain (NTD-CHC)(15). We here confirm that the second predicted Clathrin binding site (CBM2) of Ent5 binds to the *S. cerevisiae* NTD-CHC by solving the crystal structure of the complex. The CBM2 peptide engages simultaneously with the three canonical binding sites: Clathrin box, Arrestin box, and W-box (Supplementary Fig 8, see Supplementary Table 4 for Statistics). The overall backbone binding mode is conserved between CBM1 and CBM2, yet CBM2 presents distinct side-chain interactions. In the Clathrin box site (Asn89), the bulky Trp residue at position +4 cannot occupy the Leu+4 pocket as in CBM1; instead, it stacks against Phe91 via π-π interaction and forms a hydrogen bond between its indole nitrogen and Asp-1, likely stabilizing its conformation. In the Arrestin box, Trp+4 occupies the same site as Leu+4 in CBM1, enabled by a repositioning of Arg235, which also allows a cation–π interaction between Arg235 and Trp+4. In the W-box site, CBM2’s Trp+4 forms an additional hydrogen bond between its indole ring and Gln175, an interaction absent in CBM1, replacing an internal Leu+4-Leu+1-Pro-2 contact.

Proteins can be classified as *drivers* or *clients* in LLPS depending on their role in initiating phase separation and recruiting other partners. Since Ent5 has two active CBMs and can undergo phase separation under multiple *in vitro* conditions, we hypothesize that it acts as a driver of LLPS. We have then tested Ent5’s capacity for recruiting *clients* (see Fig 4). We performed recruitment experiments following Ent5-mEGFP in the presence of NTD-CHC-mScarlet and we observed that the second one is selectively recruited to Ent5 droplets (see Fig 4b). In addition, the concentration of Ent5 decreases in the presence of NTD-CHC. We have then performed the same experiment introducing mutations to the Clathrin, Arrestin and W-Box boxes of NTD-CHC (CHC CAW mutant, see Material and Methods). Ent5 cannot bind NTD-CHC CAW through their CBM1 and CBM2 regions and therefore NTD-CHC CAW is not recruited to the droplets (see Fig 4c). Our results suggest that NTD-CHC is recruited into Ent5 droplets by a well-defined, high-affinity interaction between the NTD-CHC and the CBMs in Ent5, yielding strong and efficient partitioning.

**Figure 4.**
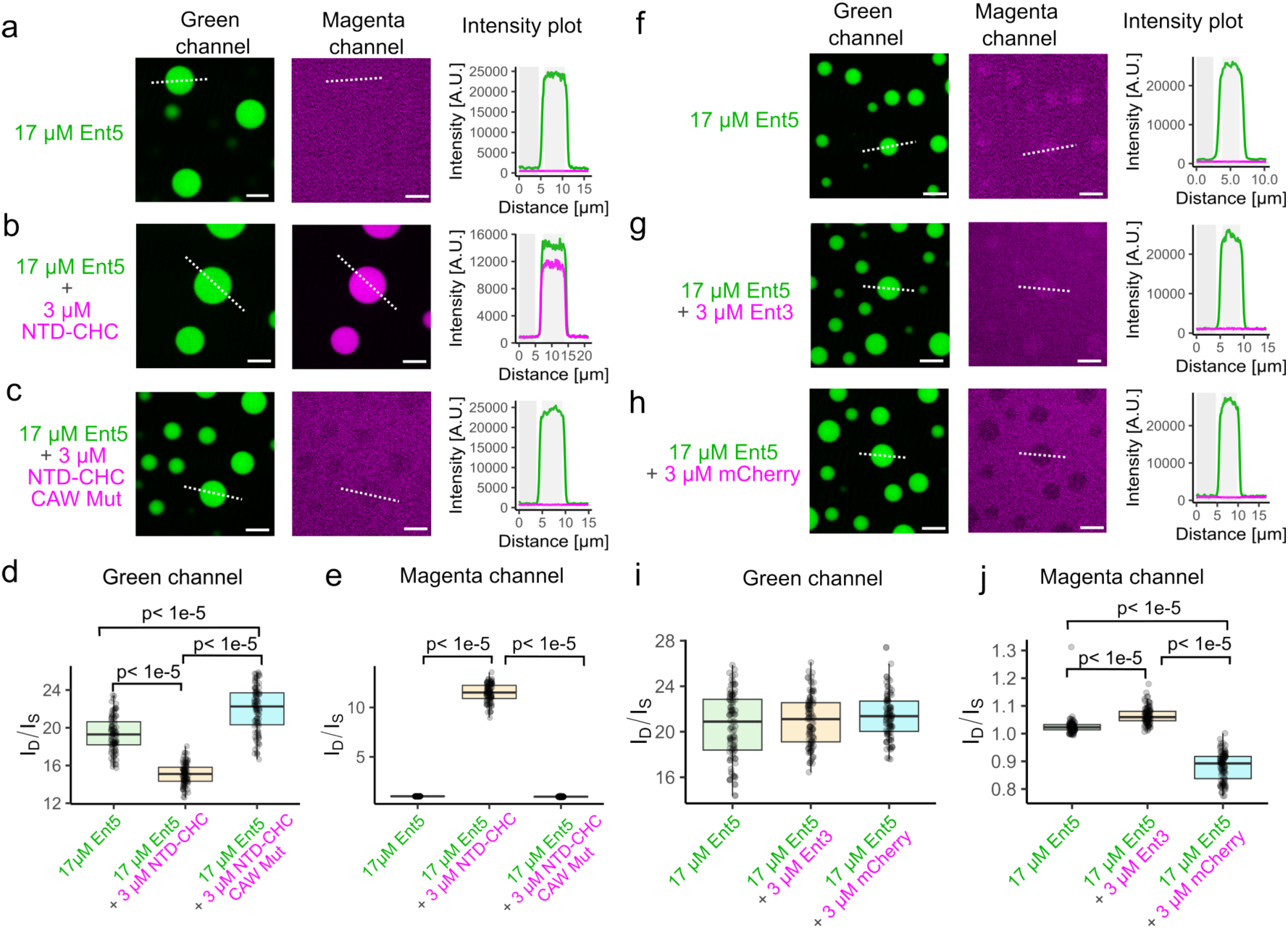
Recruitment of Clathrin Heavy Chain and Ent3 to Ent5 Droplets. a to e) Recruitment of Clathrin Heavy Chain. a) 17 μM Ent5 imaged in the green channel (488 nm excitation) phase separates without any detectable signal in the magenta channel (561 nm excitation). b) 17 μM Ent5 in the presence of 3 μM NTD-CHC-mScarlet. c) 17 μM Ent5 in the presence of 3 μM of NTD-CHC CAW mutant (Clathrin, Arrestin and W-Box mutant, incapable of binding CBM1 and CBM2 regions from Ent5, see Methods). In each microscopy panel, a representative image (n = 30) is shown with a central line drawn to quantify the difference in protein concentration between the droplet (ID, intensity in the droplet) and the surrounding solvent (IS, intensity in the solvent). (d and e) Intensity ratios (ID/IS) were quantified for the green channel (Ent5; panel d) and the magenta channel (NTD-CHC; panel e) under three conditions: 17 μM Ent5-mEGFP, 17 μM Ent5-mEGFP with 3 μM NTD-CHC-mScarlet, and 17 μM Ent5-mEGFP with 3 μM NTD-CHC CAW. Statistical tests were performed using two-sided Yuen trimmed means, corrected for multiple comparisons by the Holm–Bonferroni method; “p-holm-adj” denotes the corrected p-value. (f to j) Recruitment of Ent3. f) 17 μM Ent5 imaged alone does not show signal in the magenta channel. g) 17 μM Ent5 in the presence of 3 μM Ent3-mCherry, h) A control experiment showing 17 μM Ent5 with 3 μM mCherry shows no increase in the magenta signal is detected. (i, j) Intensity ratios (ID/IS) were quantified for the green channel (Ent5-mEGFP; panel i) and the magenta channel (Ent3-mCherry or mCherry; panel j) under three conditions: 17 μM Ent5-mEGFP alone, 17 μM Ent5-mEGFP with 3 μM Ent3-mCherry, and 17 μM Ent5-mEGFP with 3 μM mCherry. Statistical tests were conducted using two-sided Yuen trimmed means, corrected for multiple comparisons by the Holm-Bonferroni method. White scale bars in all microscopy panels represent 5 μm.

Ent3 is an epsin involved in the recruitment of Gga and AP-1 proteins during trans-Golgi network trafficking(26) and does not contain a Clathrin binding motif. In order to test Ent5 capacity as a LLPS driver, we performed fluorescence microscopy colocalization experiments with Ent5-mEGFP and Ent3-mCherry to test the recruitment of Ent3 to Ent5 droplets (see Fig 4 f-j and Supplementary Fig 9). Our results indicate that Ent5 increases Ent3 concentration in the droplet phase (Fig 4j) but Ent3 does not change the concentration of Ent5 (Fig 4i). Actually, the recruitment of Ent3 to Ent5 droplets depends on the Ent5 concentration which is consistent with Ent3 being a LLPS *client* (Supplementary Fig S9a). As a control, we show that mCherry alone does not colocalize within Ent5 droplets, meaning it does not behave as a LLPS *client* (Supplementary Fig. S8b). In contrast, to what has been observed for NTD-CHC, Ent3 partitions into Ent5 droplets but does so via an unknown mechanism.

### Sla2 behaves both as an LLPS *driver* and *client*

Through the same analysis as done on Ent5 with *client* proteins, we have performed Sla2 experiments to test the interaction with protein partners on the droplet phase. Ede1, homologous to *Hs*Eps15, has been described as a *driver* in phase separation and its central region, residues 366 - 900 (Ede1 IDR-CC), has been implicated in the formation of condensates at ∼5 µM protein concentration(27). We therefore tested the co-localization of Ede1 IDR-CC and Sla2 IDR-CC (see Fig 5a-d).

**Figure 5.**
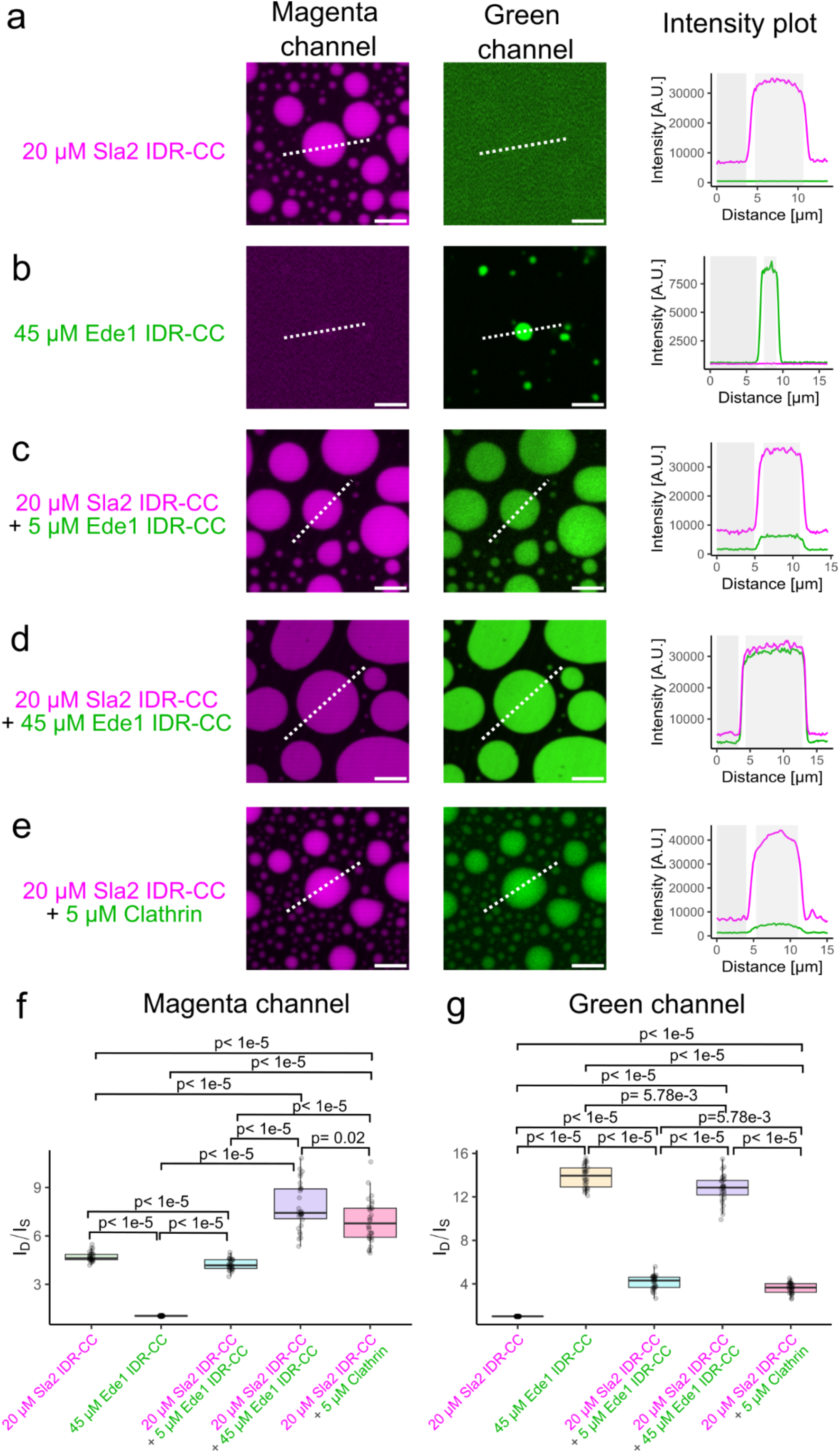
Co-localization and Recruitment of Sla2 with Ede1 and Clathrin in Condensates. a to e) Representative droplets for each sample are shown for the magenta (left panels) and the green (middle panels) channels. Measurements across the white dotted lines are used to quantify fluorescence intensities between the droplet (ID) and the surrounding solvent (IS). Intensity profiles are plotted on the right panels.. Images corresponding to Sla2 IDR-CC-mScarlet. a,c, d and e) and for Ede1-mEGFP (b,c and d) or Clathrin-mEGFP e). f) Average intensity ratios (ID/IS) measured from the magrenta channel for n = 30 droplets. Statistical tests were conducted using two-sided Yuen trimmed means, corrected for multiple comparisons by the Holm–Bonferroni method; “p-holm-adj” denotes the corrected p-value. g) Average intensity ratios (ID/IS) measured from the green channel for n = 30 droplets. Statistical tests were conducted using two-sided Yuen trimmed means, corrected for multiple comparisons by the Holm-Bonferroni method; “p”. White scale bars in all microscopy panels represent 5 µm.

The ID/IS ratio for Sla2 IDR-CC at 20 µM is 4.7, and the distribution of protein is uniform throughout the droplet. When mixing it with an excess of Ede1 IDR-CC, 45 µM, Sla2 ID/IS is 7.8 and Ede1 ID/IS is 12.8 (see Fig 5f and g). An indication that Ede1 increases the condensation of Sla2 into droplets. In this case Ede1 is the *driver* and Sla2 a *client*.

It has been shown that Sla2 binds Clathrin Light Chain (CLC) through two binding sites(16). Utilizing the full-length CLC and CHC distal leg (residues 1172-1572) heterodimer construct conjugated to mNeonGreen, we have measured the recruitment of Clathrin to Sla2 IDR-CC–mScarlet droplets (see Fig 5e). The ID/IS of Sla2 is significantly increased from 4.7 to 6.9 (Fig 5f) and the ID/IS of Clathrin from 1.01 to 3.6 (Fig 5g), indicating a recruitment of Clathrin to Sla2 droplets.

### Ent5 and Ent1/Sla2 condensation is promoted by the membrane environment

We have used Giant Unilamellar Vesicles (GUVs) as a vehicle for determining the recruitment of both Ent5 and Sla2 condensates to membrane environments (see Fig 6). In the case of Ent5 the ENTH domain binds PI(3,5)P2(18) and therefore the GUVs were prepared with 8% PI(3,5)P2 and 0.1% Cy5.5-labelled PE (1,2-distearoyl-sn-glycero-3-phosphoethanolamine-N) to visualize the membrane in fluorescence microscopy. 5% PEG 8000 was chosen for the condensate-inducing variable as salt modification is not possible due to destroying the GUVs when changing the molality of the external solution as opposed to the internal sucrose solution. Our results show that Ent5 forms localised larger patches corresponding to condensates attached to the membrane at 3.5 µM protein concentration in the presence of the crowding agent (see Fig 6b). This phenotype is not observed for similar concentrations of Ent5 ΔHelix mutant in the presence of 5% PEG 8000 (see Fig 6c), displaying a behavior similar to the wt Ent5 in the absence of crowding agent (see Fig 6a). Therefore, our results suggest that the helix is required for condensation once Ent5 is bound to the membrane.

**Figure 6.**
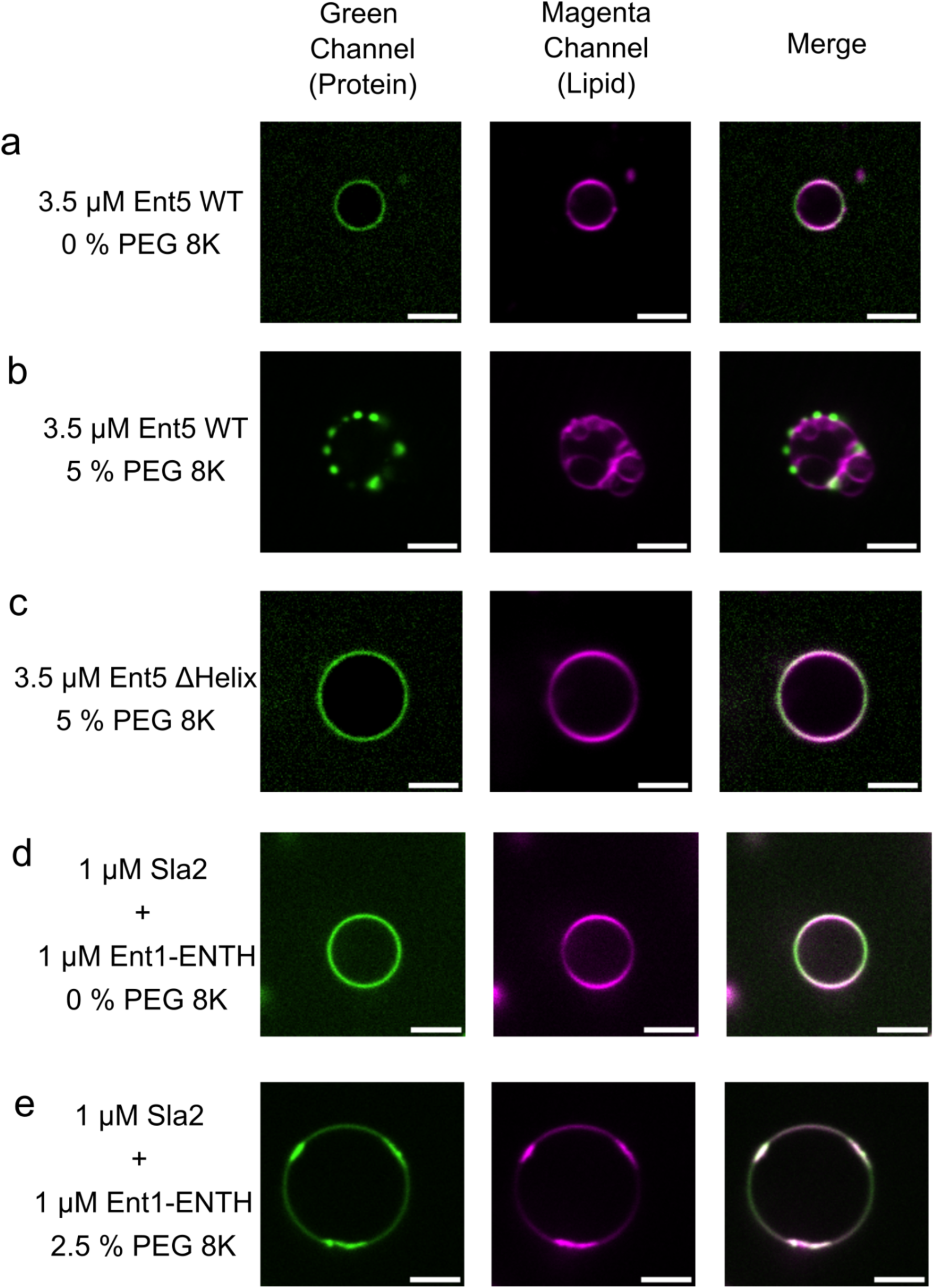
Protein Condensates are recruited onto GUVs Containing Signaling Lipids. a-c) Preformed GUVs, containing 8% PI(3,5)P2, were incubated at 21 °C with 3.5 μM Ent5-mEGFP. Fluorescence images were acquired in two channels: the protein is visualized in the green channel (mEGFP) and the lipid in the magenta channel (Cy5.5). Three conditions are shown: a) 0% (w/v) PEG 8000, b) 5% (w/v) PEG 8000 and c) 3.5 μM Ent5 ΔHelix-mEGFP in the presence of 5% (w/v) PEG 8000. d-f) Preformed GUVs containing 8% PI(4,5)P2 were incubated at 21 °C with a mixture of 1 μM Sla2–mEGFP and 1 μM Ent1-ENTH. Two conditions are displayed: d) 0% (w/v) PEG 8000 and e) 2.5% (w/v) PEG 8000. White scale bars in all images represent 5 μm.

Sla2 binds PI(4,5)P₂ when forming a complex with the epsins Ent1 and Ent2(28) through their ANTH and ENTH domains, respectively, forming a high-affinity complex in the nanomolar range(29) that induces membrane deformation(30). In order to test the formation of condensates onto the membrane environment, GUVs were prepared containing 8% PI(4,5)P2 and 0.1% Cy5.5. We mixed full-length Sla2-mEGFP with equimolar amounts of Ent1 ENTH domain at 1µM protein concentration in the GUV mix. This working concentration is below the concentration of protein needed to trigger membrane deformation, which has been observed to be higher than 10 µM for full-length Sla2 (data not shown). In our experiments, we observe uniform recruitment of Sla2-mEGFP to the membrane but no clustering (see Fig. 6d). In the presence of crowding agents local condensates of increased protein concentration are seen but no tubular deformation of the membrane (see Fig. 6e).

### Ent5 Helix removal prolongs Lifetime of Gga2-depedent trafficking

We found that a dynamic helix (residues 249-287) of Ent5 behaves as a *sticker* for LLPS *in vitro*. To test its functional relevance *in vivo*, we generated a construct lacking this helix (ΔHelix) and asked whether its removal alters adaptor-mediated trafficking. To read out trafficking dynamics, we quantified the lifetimes of Gga2 and Ent5 puncta together as a proxy. For this, we developed a dual-camera, 3D fluorescence microscopy assay in which endogenous Gga2 is C-terminally tagged with mNeonGreen, and Ent5 is supplied from a plasmid as either WT or ΔHelix, tagged with mScarlet (see Fig. 7).

**Figure 7.**
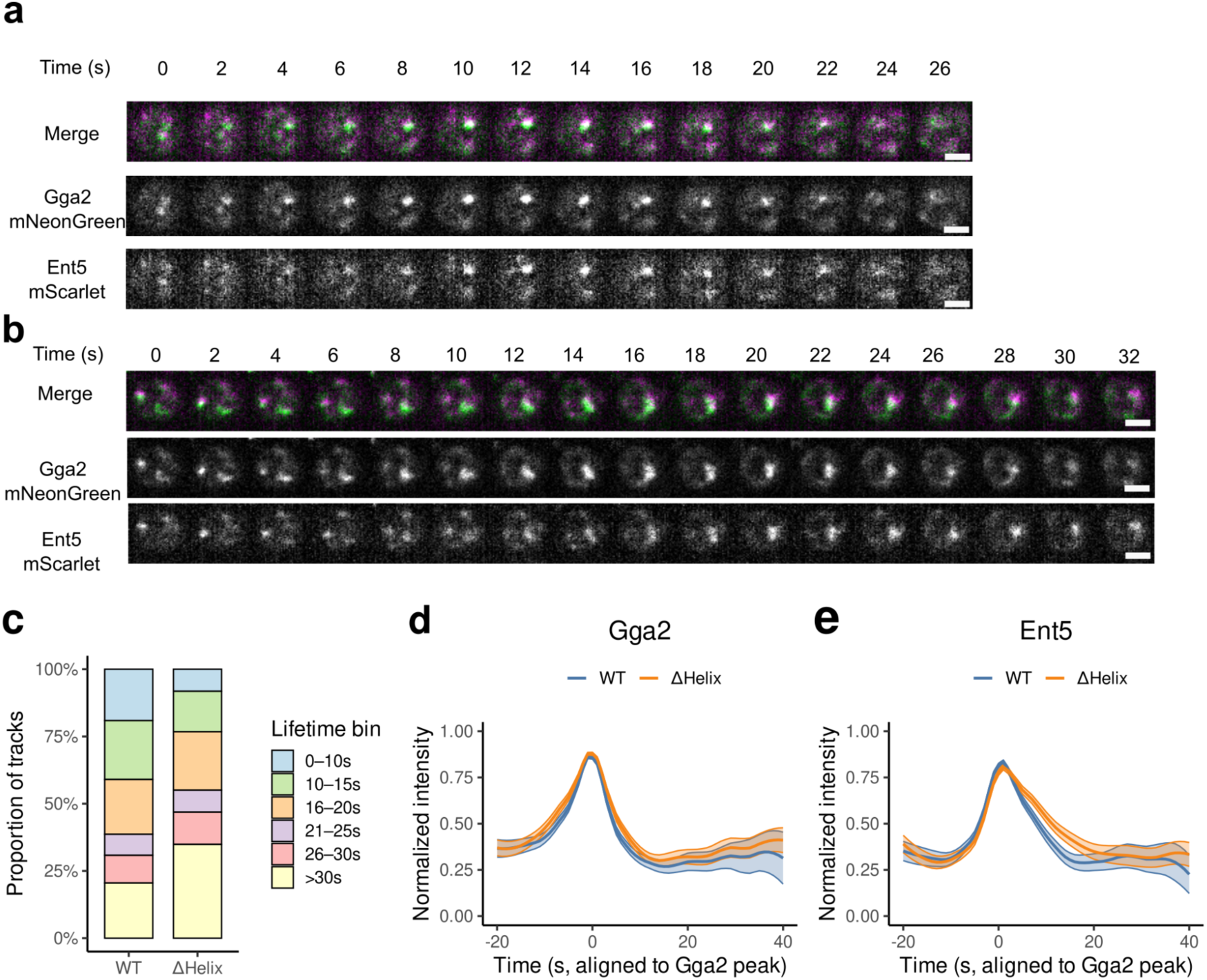
Removal of the Ent5 helix prolongs Gga2-associated trafficking events. a–b) Representative time-lapse montages centered on tracked spots for WT a) and ΔHelix. b) Gga2 (Green), Ent5 (Magenta); merged composites shown. Scale bar, 2 µm. c) Proportional stacked bar plots showing the distribution of track lifetimes across six duration bins (0–10, 10–15, 16–20, 21–25, 26–30, >30 s) for WT (left) and ΔHelix (right). Statistical comparison using a χ² test of independence on the full contingency table: χ² = 54.43, df = 5, p = 1.71 × 10⁻^10^. d–e) Mean normalized intensity profiles of Gga2 d) and Ent5 e) for WT (blue) and ΔHelix (orange). Traces are aligned to the Gga2 intensity peak (t = 0), with each curve showing the LOESS fit to the dataset mean (solid line) and 95% confidence interval (shaded region). Individual trajectories were not smoothed prior to averaging. Sample sizes: WT, n = 639; ΔHelix, n = 525.

Across movies, the Ent5 ΔHelix mutant prolongs Gga2-positive events. This shift in the average reflects a redistribution of event durations when all tracks are pooled: the fraction of short events (0–10 s) becomes less frequent, while long events (>30 s) increase in ΔHelix relative to WT (Fig. 7c; χ² = 54.43, df = 5, p = 1.71 × 10⁻¹⁰; nWT = 639, nΔHelix = 525; examples: 0–10 s WT 19.1% → ΔHelix 8.2%; >30 s WT 20.5% → ΔHelix 34.9%). Peak-aligned overlays of Gga2 (LOESS fit shown as a solid line, 95% CI as shaded area) reveal similar assembly–disassembly kinetics in both genotypes (Fig. 7d).

For Ent5 dynamics, the ΔHelix mutant also increases the average event lifetime (Fig. 7e; LOESS overlays, nWT = 639, nΔHelix = 525), while representative montages show comparable event morphology in both conditions (Fig. 7a–b). Together, these results indicate that removal of the Ent5 helix selectively extends the duration of Gga2-marked trafficking events and shifts the population toward longer-lived assemblies, without altering Gga2 kinetics but with a detectable change in Ent5 recruitment dynamics.

## Discussion

Our experimental screening revealed that predictions for phase separation can be misleading. These predictions are based primarily on the presence of intrinsically IDRs to suggest LLPS. Despite computational predictors suggesting a potential role in LLPS, we did not observe condensate formation for Ent1, Ent2, or Ent3. Consistently, these epsins showed no signs of phase separation in our chemical screening assays. Supporting this, Sarkar et al. (2025)(31) reported that Epsin1 not only fails to form condensates but may also destabilize the condensation of Eps15.

Another observation is that predictions for phase separation are sequence-based, with algorithms taking into account the local residue context but not structural information. Therefore, while the NTD-CHC domain is not predicted to undergo LLPS, the CLC, which contains an IDR, is predicted to spontaneously undergo LLPS. However, we do not know how the assembled triskelions would behave *in vivo*. Adaptors bound to them seem to act as more intuitive switches for condensate formation.

Ent5 demonstrated clear phase separation in our chemical screening, requiring both its helical region and the ENTH domain. Notably, the full IDR complements at a lower concentration than the isolated helix, indicating that the larger IDR, acting as a *spacer,* contributes significantly to phase separation. Our findings suggest that Ent5 may act as a driver of phase separation in endosomal trafficking given its capacity for recruiting NTD-CHC and, to a lesser degree, Ent3.

Sla2 appears to act as a client of Ede1, which arrives earlier in the endocytic pathway. Ede1 enhances Sla2 condensation, highlighting a potential temporal regulation of condensate formation(14). However, we show here that individual regions of Sla2 are capable of phase separation. The central (CC) region behaves as a *sticker*, and complementation experiments indicate inter-protein interactions that are further enhanced in the context of the full-length protein. This suggests a dual client/driver role for Sla2, potentially tied to its critical function in regulating the Pan1/End3/Sla1 complex (32, 33) . If this role extends to other adaptors and regulators, it may facilitate the recruitment of actin polymerization and regulatory machinery during endocytosis.

Our results further underscore the importance of discrete structural elements in driving phase separation. In Ent5, we identified a dynamic helix within the IDR that functions as a bona fide *sticker.* This helix alone is insufficient to promote condensation, but together with the surrounding disordered region and the ENTH domain, it enables robust LLPS at physiological concentrations. The synergy between inducible secondary structure and flexible spacer regions appears to be a recurring principle in endocytic adaptors. In Sla2, by contrast, the central CC domain fulfills the role of a *sticker*, mediating intermolecular interactions that drive condensation. Unlike the isolated helix of Ent5, the CC-region of Sla2 is sufficient to form droplets under crowding conditions, and its condensation is further strengthened when combined with adjacent domains. These findings highlight that endocytic adaptors exploit distinct structural motifs as determinants of LLPS.

When analysing condensate formation in the context of lipidic membranes, both the ENTH domain in Ent5 and the ANTH domain in Sla2 synergistically facilitated LLPS. Additionally, we observed that phosphoinositides (PIPs) help concentrate adaptor proteins on membranes, thereby promoting the nucleation of protein condensates. This protein clustering could be beneficial in facilitating trafficking processes, as it may locally increase adaptor density at the membrane and favor efficient cargo recruitment.

The functional role of these sequence features is further supported by our *in vivo* experiments. Deleting the dynamic helix of Ent5 impaired its ability to form condensates on membranes and altered adaptor-mediated trafficking. Using total lifetime as a readout, we observed that ΔHelix prolonged trafficking events and shifted the population from predominantly short-lived to longer-lived tracks. Importantly, this change in lifetimes occurred without detectable alterations in the recruitment sequence of endocytic proteins, suggesting that condensate dynamics influence the timing of adaptor turnover rather than the order of recruitment. These data provide direct evidence that the structural determinants of LLPS uncovered *in vitro* also modulate trafficking dynamics in living cells. Consistent with this, Hung and Duncan (2016) (24) reported that disrupting Ent5’s clathrin-binding function impairs coat maturation and prolongs the lifetime of clathrin-coated structures. In their study, mutation of the clathrin-box motifs or complete deletion of *ENT5* led to significantly extended lifetimes of Gga2-marked endosomal/TGN coats, indicating delayed coat disassembly. Our ΔHelix mutant produces a similar phenotype: endocytic trafficking events persist longer than normal (shifting from mostly short-lived to predominantly longer-lived tracks) despite an unaltered recruitment sequence. This parallel suggests that both the clathrin-binding motifs and the phase-separating helix of Ent5 are critical for timely coat turnover

Since clathrin-adaptor interactions are inherently weak in binding affinity(15, 34–36), condensation could contribute to selective recruitment. Not all proteins can partition into a given condensate. Instead, phase-separated condensates concentrate specific client molecules while excluding others(37–39). This selectivity suggests that drivers of condensates act as biochemical filters that preferentially recruit key functional proteins.

In summary, our study shows that distinct structural features in clathrin adaptors act as determinants of LLPS, enabling these proteins to function as drivers or clients depending on context. Together with their ability to engage clathrin and membranes, phase separation emerges as a shared strategy by which adaptors regulate endocytosis and trafficking.

## Material and Methods

### Recombinant protein expression

All constructs were expressed in *E. coli* as follows. Wild-type and variant Ent5 and Ent3 fusions (6×His-Sumo3-Ent5-mEGFP, 6×His-Sumo3-Ent3-mCherry, mCherry, plus all Ent5-mEGFP variants) were cloned into pETM-11 (N-terminal 6×His-Sumo3 tag). The N-terminal domain of yeast clathrin heavy chain (NTD-CHC, 1-369) and Ent1-ENTH/CLC were cloned into pETM-30 (N-terminal His-GST tag with TEV site). Full-length Sla2 and variants, Ede1 (366–900), and CHC (1172-1574) were cloned into pETM-11 and co-expressed with pLysS.

Plasmids were transformed into chemically competent *E. coli* Rosetta2 BL21(DE3) (Ent5/Ent3, Ent1/CLC), BL21(DE3) (NTD-CHC) or BL21(DE3)pLysS (Sla2/Ede1/CHC).

Transformants were selected on LB agar with 30 µg/mL kanamycin (all) and 34 µg/mL chloramphenicol (Rosetta2) at 37 °C overnight. Several colonies were inoculated in an LB starter culture (same antibiotics) at 37 °C, 200 rpm overnight.

The overnight culture was diluted 1:100 into fresh medium (Ent5/Ent3: LB; ScChc-NTD: 2×TY [16 g tryptone, 10 g yeast extract, 5 g NaCl per L]; Sla2/Ede1/CHC: Terrific Broth) and grown at 37 °C, 200 rpm until OD₆₀₀ reached 0.8–1.0 (for Sla2/Ede1/CHC: 0.6–0.8). Cultures were then cooled: to 18 °C for Ent5/Ent3, ScChc-NTD and Ent1/CLC; to 16 °C for Sla2/Ede1/CHC. Protein expression was induced with IPTG (0.3 mM for ScChc-NTD; 1 mM for all others) and allowed to proceed 14-18 h at the induction temperature with shaking.

Cells were harvested by centrifugation for 15 min at induction temperature,7,000 × g for Ent5/Ent3, Sla2/Ede1/CHC; 12,000 × g for ScChc-NTD,and pellets were stored at -20 °C until purification.

### Recombinant Protein Purification

Recombinant protein purification was performed at 4 °C unless otherwise noted. Frozen cell pellets were thawed on ice and resuspended in lysis buffer (5-6 mL per gram of pellet; see Supplementary Table S2). The suspension was rotated at 8 °C until all visible clumps had dispersed, then lysed by three to four passes through an Emulsiflex C3 homogenizer (Avestin) at 15 kPsi. The crude lysate was clarified by centrifugation at 43,700 ×*g* for 1 h at 4 °C and subsequently filtered through a 0.45 µm membrane.

For His-tagged Ent5, Ent3 and the N-terminal clathrin heavy chain domain (NTD-CHC), the cleared supernatant was loaded at 2 mL/min onto two serial 5 mL HisTrap HP columns (GE Healthcare) pre-equilibrated in buffer B1 followed by buffer A1 (see Table S2). After washing with 20 column volumes of buffer A1 and 5 CV of 5% buffer B1, bound protein was eluted over a 10 CV linear gradient to 100% buffer B1, and 1.5 mL fractions were collected. Sla2, Ede1 and CHC^1172-1574^ constructs were purified by batch binding to gravity-flow Roti®Garose Ni–NTA resin (Carl Roth), which was equilibrated in buffer A2, washed with 10 CV of (90% Buffer A2: 10% Buffer B2), and eluted with 10 CV of buffer B2; 1 mL fractions were collected throughout. Fractions containing target proteins were identified by A₂₈₀ nm absorbance and SDS-PAGE, then pooled for tag removal.

His-tagged samples (Ent5, Ent3, NTD-CHC, Sla2, Ede1 and CHC^1172-1574^) were dialyzed overnight at 4 °C against Buffer 1 (for Ent5, Ent3 and NTD-CHC) or Buffer 2 (for the other constructs) in the presence of SenP2 SUMO protease at 1 mg per 25 mg of fusion protein. NTD-CHC was cleaved instead with TEV protease at the same enzyme: substrate ratio. GST-tagged Ent1-ENTH and CLC were bound to Glutathione Sepharose 4B (Cytiva), washed in buffer A2, and eluted by on-column digestion with 1 mg TEV per mL resin in buffer A2 overnight at 4 °C.

To remove protease and residual contaminants from Ent5 and Ent3, dialyzed samples were adjusted to 0.7 M ammonium sulfate and applied at 4 mL/min to a HiTrap Butyl HP column (GE Healthcare) pre-equilibrated in Milli-Q water, then HIC buffer A. After a 10 CV wash in HIC buffer A, proteins were eluted over an 8 CV gradient to 35% HIC buffer B, held at 35% for 5 CV, and ramped to 100% over 5 CV. One-milliliter fractions were pooled based on SDS–PAGE, concentrated to ∼2 mL (Amicon Ultra-15, 10 kDa cutoff), and cleared of precipitate by centrifugation at 21,000 × g for 20 min at 4 °C. All other cleaved His-tag constructs (Sla2, Ede1, Ent1-ENTH, CLC, CHC^1172-1574^) were passed three times over gravity-flow Ni-NTA resin equilibrated in their dialysis buffer; the combined flow-through and a 5% imidazole wash were pooled, concentrated, and clarified by centrifugation at 21,000 × g for 20 min at 4 °C.

Final polishing was achieved by size-exclusion chromatography. Ent5 and Ent3 were run on a HiLoad 16/600 Superdex 200 pg column equilibrated in SEC Buffer 1, NTD-CHC on Superdex 200 pg in SEC Buffer 2, and Sla2, Ede1, Ent1-ENTH, CLC, and CHC^1172-1574^ on Superose 6 (Sla2/Ede1, CHC) or Superdex 200 pg (Ent1-ENTH and CLC) in SEC Buffer 3 (all buffers in Table S2). Samples (≤2 mL) were injected and eluted at 0.8 mL/min, and 1 mL fractions corresponding to the main peak,verified by SDS-PAGE,were pooled, concentrated, and cleared of aggregates by centrifugation at 21,000 × g for 20 min at 4 °C. Purified proteins were aliquoted, flash-frozen in liquid nitrogen, and stored at -70 °C.

### Construct Details and Site-Directed Mutagenesis

Mutations in the plasmids pETM-30-NTD-CHC and pETM-11-Ent5-mEGFP were introduced using a modified QuickChange protocol. Overlapping mutagenic primers (see Table S3) were designed to amplify the entire plasmid. Each 50 µL PCR reaction contained 100 ng template DNA, 200 nM of each primer, and 25 µL of 2× Phusion reaction mix [40 mM Tris–HCl, pH 8.8 (25 °C); 4 mM MgCl₂; 120 mM KCl; 20 mM

(NH₄)₂SO₄; 0.02 mM EDTA; 0.2% Triton X-100; 8% glycerol; 0.005% Xylene Cyanol FF; 0.05% Orange G; 0.4 mM dNTPs; and 0.04 U/µL Phu-Sso7d polymerase with nuclease-free water. The PCR cycling conditions were as follows: initial denaturation at 98 °C for 3 min; 20 cycles of 98 °C for 1 min, 56 °C for 45 s, and 72 °C for 7 min; followed by a final extension at 72 °C for 10 min. After amplification, 1 µL of DpnI was added, and the reaction was incubated at 37 °C for 2 h to digest the methylated parental DNA. A 10-µL aliquot was then transformed into chemically competent *E. coli* DH5α cells. The desired mutations were confirmed by Sanger sequencing (Microsynth, Göttingen, Germany). The CAW Clathrin mutant was designed as follows: for the Clathrin box, we introduced mutations K63E, I87D, Q89A and K98E; for the Arrestin box, Q195A, I197T and K251E; and for the W-Box, we mutated residues F26A and Q155A.

### Peptide synthesis

Peptides were purchased from NovoProLabs (Shanghai, China) with a purity of at least 98% and TFA-free (Acetate-salt). Ent5 Helix(249-287, ANSNTRRRSHMEEQRRQRREILREQIKNKEQQRKRKQQ) and Ent5-6P (249-287, ANSNTPRRSHMPEQRRPRREPLREQIPNKEQQPKRKQQ). Peptide stocks were solubilized in water at a concentration of 20 mM and then diluted again at a final working concentration in 10 mM Tris, pH 7.5.

### GUV production and imaging

Using Platinum nets dry 20 µL lipid mixture at a final concentration of 1.25 mg/ml. Mixture contains (all molar %) 76% DOPC, 0.1% DOPE, 16% DOPS, and 8% PI(3,5)P2 for GUVs binding Ent5 and 8% PI(4,5)P2 for GUVs used in experiments involving Sla2. Spread on both nets three times, drying for at least 1h in a fume hood to remove residual CHCl3. A continually heated sucrose solution at > 40℃ matching the molality of the SEC Buffer is added to the cuvette to submerge the Pt nets for 30 minutes. Apply an AC for 2 h, 2.2 V at 10 Hz, then for 20 min apply 2.2 V at 2 Hz.

For imaging, the slide area is incubated with 5% BSA in PBS for 1h, washed five times with PBS and left soaked in PBS. Add the protein-PEG solution to the slide and subsequently add a one-in-twenty dilution of GUV sucrose solution. Imaging was done using a Nikon Ti2 spinning disk microscope using the 488 nm and 640 nm excitation wavelengths.

### Mass Photometry (MP)

MP experiments were performed using a Refeyn TwoMP mass photometer with AcquireMP software (Version 2024 R1). Protein samples of Ent5WT-mEGFP were adjusted to 1 µM in Experiment Buffer 100 (composition in Supplementary Table S2). The instrument was auto-focused using 18 µL of Experiment Buffer 100 or LLPS Trigger Buffer (see Supplementary Table S2 for composition). Next, 1-2 µL of the protein stock was applied to the measurement chamber and recorded for 1 min at room temperature. Calibration was carried out using the NativeMarker Unstained Protein Standard (Invitrogen, LC0725). Data analysis was performed with DiscoverMP and further processed using custom Python scripts (https://github.com/steniegit/BioPhyPy) based on the eSPC (http://www.spc.embl-hamburg.de) framework (Niebling et al., 2022).

### Fluorescence Microscopy and FRAP

Images and time-lapse videos of protein droplets were acquired on a Nikon Eclipse Ti2 microscope equipped with a spinning disk confocal unit, an oil-immersion 100× objective (NA 1.49), and an iXon Ultra 888 EMCCD camera. For imaging, mEGFP was excited at 488 nm (1% laser power) and using a bandpass emission filter 525/50, mCherry was excited at 561 nm (1% laser power), and mScarlet at 561 nm (10% laser power) and a bandpass emission filter 600/25 was used for imaging. For photobleaching (FRAP) experiments, 473 nm and 522 nm lasers were used at 40% power (Rapp OptoElectronic GmbH, Wedel, Germany). Movies were recorded at 500 ms intervals for 90–180 s, with the acquisition starting 10 s before bleaching to establish a baseline.

Samples were prepared by dialyzing proteins overnight in a buffer containing 150 mM Tris (pH 7.5), 100 mM NaCl, and 0.5 mM TCEP (see Supplementary Table S2). Immediately before imaging, samples were diluted 1:5 in a Tris buffer (pH 7.5, 0.5 mM TCEP) to induce LLPS, resulting in a final buffer composition of approximately 100 mM Tris, 20 mM NaCl, and 0.5 mM TCEP. In experiments where salt-induced LLPS was insufficient, 5 % (w/v) PEG 8000 was added as an alternative trigger.

For quantitative analysis, 99 droplets per condition were examined to determine the partition coefficient (ratio of droplet intensity, *I*D, to the surrounding solution intensity, *I*S).

Statistical significance was assessed using the Mann-Whitney U test. FRAP recovery curves were corrected for photobleaching and normalized using custom Python scripts. The recovery kinetics were fitted to the equation

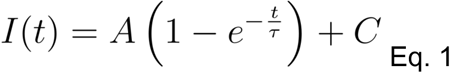

where τ is the recovery time constant, *A* is the mobile fraction, and *C* is the baseline offset. The half-time of recovery (*t*1/2) was calculated according to established methods (40). The fitting parameters of Ent5 FRAP curves are A = 0.45, tau = 19.2s and C = 0.43 while for Sla2 are A = 0.47, tau = 112.8 s, C = 0.3

### Circular Dichroism Spectroscopy

Peptides corresponding to the WT and 6P mutants were purchased from NovoProLabs (Shanghai, China) at ≥98% purity. Trifluoroacetic acid (TFA) was removed and exchanged with acetic acid before use. CD spectra were recorded on a Chirascan spectrophotometer by averaging three scans at a bandwidth of 1 nm. Measurements were performed in either 1 mm or 5 mm high-precision cuvettes (Hellma Analytics) at a peptide concentration of 20 µM in 10 mM Tris (pH 7.5). The sample temperature was controlled at 20 °C using a TC125 temperature controller (Quantum Northwest, Liberty Lake, USA) unless otherwise stated. Helical content was assessed via a TFE titration series from 0% to 52% (v/v). The free energy of helix formation was determined by fitting the data to a two-state model (Equation [2]) using custom SPC ChiraKit (41).

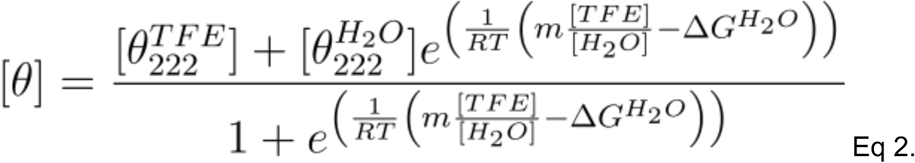

### Stopped-flow Light Scattering

Stopped-flow light scattering measurements were performed on a µSFM stopped-flow (BioLogic, Grenoble, France) fitted with a MOS-200/M spectrometer. Syringe 1 (100 mM Tris pH 7.5, 0.5 mM TCEP) and syringe 2 (18 mg/mL Ent5-WT in 100 mM Tris pH 7.5,

0.5 mM TCEP, 150 mM NaCl) were rapidly mixed (9:1 buffer:protein, 32 µL total volume) at ∼1 mL/s into a µFC-08 cuvette maintained at 8 degrees. Measurements were done at 600 nm, detected at 144 V, and sampled at 0.5 ms intervals over 4001 points (0–2 s). Data acquisition began immediately upon mixing. All kinetic traces were fitted in R using non-linear least squares using two exponential curves using the following equation:

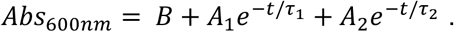

where B, A1 and A2 are pre-exponential constants, t is the time in milliseconds and τ_%_ and τ_*_ are the two time constants of the process.

### Stopped-flow time-resolved SAXS

Time-resolved SAXS measurements were performed at the SAXS beamline P12 (EMBL, PETRA III, DESY, Germany) using a stopped-flow mixer (µSFM, BioLogic, Grenoble, France) equipped with a quartz capillary (1.0 mm inner diameter) as a sample delivery system (42). The device was used to rapidly mix 17.6 mg/ml Ent5-WT in high-salt buffer (100 mM Tris pH 7.5, 0.5 mM TCEP, 150 mM NaCl) with low-salt buffer (100 mM Tris pH 7.5, 0.5 mM TCEP) in a 1:9 ratio with a total volume of 32 µl. Measurements were done at 8 degrees keeping the temperature constant with a circulation water bath (Huber - Ministat 240 Thermocycler (Offenburg, Germany)).

SAXS curves were recorded with an Eiger X 4M detector positioned at a distance of 3 m from the sample using delays ranging from 0 to 380 ms in 20 ms increments, with an exposure period of 20 ms per frame. Two independent series were acquired and averaged for each time point within a q-range of 0.02 and 3.0 nm^-1^. To visualize the time evolution of the reaction, the intensities at q = 0.26 nm^-1^ were plotted against the reaction time.

### Protein crystallization and data collection

Purified Ent5 concentrated up to 10 mg/ml was mixed with the Ent5.2 peptide (IDDLLDWDGP, NovoProLabs, China) at a final concentration of 8.8 mM, and the mixture was set for co-crystallization using commercially available screens. After one day of incubation, a crystal appeared in a condition consisting of 0.056 M sodium phosphate and 1.334 M potassium phosphate. The crystal was retrieved from the droplet and briefly soaked in crystallization buffer saturated with Ent5.2 peptide at 8.8 mM and glycerol as cryo-protectant at 10% v/v, followed by flash-freezing under liquid nitrogen for storage and x-ray data collection. X-ray diffraction data were collected at the P14 beamline of the PETRA III synchrotron.

### X-ray diffraction data processing

Raw x-ray diffraction data indexing and integration with XDS(43) identified C121 space group followed by XSCALE scaling step with data extending to 1.7 Å resolution. Molecular replacement succeeded using the 9EX5 PDB structure as a search model and yielding a solution scoring a TFZ of 76.6 that could place two molecules in the asymmetric unit. The resulting molecular replacement solution displayed a well-defined electron density with clear additional positive density maps tenable for structure modelling. Subsequently iterative rounds of model building and structure refinement were achieved with Coot(44) and Refmac(45) .

### Phylogenetic Tree Reconstruction and AlphaFold3 Modeling

Homologs of ScEnt5 (UniProtKB Q03769; strain S288c) were identified in UniRef100 (March 2025) using BLAST+ v2.15(46) (blastp; E-value ≤ 1e-5; soft masking on), then used to build an Ent5-specific profile with MUSCLE v5.1(47)(seed MSA) and HMMER v3.4 (hmmbuild) (48), followed by iterative jackhmmer searches against UniRef100 (N = 5; incE = 1e-5; domE = 1e-5) to expand the set while limiting drift. To enforce the Ent5-specific ENTH-domain insertion, each retained sequence was modeled with AlphaFold3(49) and structurally aligned to the yeast Ent5 ENTH reference using TM-align 20160521(50); reference residues 86-98 were evaluated within a buffered window 80-103, and sequences were kept only if at least half of the reference-mapped columns were non-gaps in the candidate (≥ 50% occupancy), to check for the insertion specific to Ent5 (51). The final sequence set was aligned with MUSCLE v5.1 (47) (Super5 for large N; trimming of highly gapped/poorly aligned sites prior to inference) Lineage metadata were retrieved from NCBI Taxonomy(52) (snapshot June 2025) for downstream annotation. Maximum-likelihood trees were inferred with IQ-TREE v2.3.6(53) using ModelFinder for automatic model selection, and assessed branch support with 1,000 ultrafast bootstrap replicates refined by NNI together with 1,000 SH-aLRT replicates. lineages represented by fewer than three sequences were removed prior to visualization. Trees were rendered in iTOL v7.2.1(54) (circular layout) and exported as SVG for figure preparation in Inkscape.

### *In vivo* Gga2-Ent5 recruitment dynamics

*Saccharomyces cerevisiae* cells (MK100 WT: MATa; his3Δ200; leu2-3,112; ura3-52; lys2-801) were made competent and transformed as previously described(15) to tag Gga2 with mNeonGreen at the C-terminus. Additionally, cells were transformed with pRS315-Ent5-mScarlet (WT or ΔHelix; LEU selection marker), and the endogenous ENT5 was then knocked out (URA cassette). Yeast cells were grown overnight at 30 °C with shaking in a 24-well plate using LD (low-fluorescence SD)-Trp^-^,Leu^-^ Ura^-^ medium (yeast nitrogen base without amino acids supplemented with the corresponding dropout mix; Formedium, CYN402, with 2% Glucose). Cells were diluted into fresh medium to OD600 = 0.1 and grown at 30 °C with shaking for 4-6 h to log phase (OD600 = 0.6-1.2). 8-well glass-bottom μ-slides (Ibidi, 80807) were coated with 50 µL Concanavalin A (1 mg/mL; Sigma-Aldrich, C2010; in 10 mM sodium phosphate pH 6, 10 mM CaCl2, 1 mM MnCl2) for 5 min and washed twice with 50 µL fresh medium. Then 50 µL of cell suspension were applied for 5 min, removed, and wells were washed twice with 50 µL fresh medium to remove unattached cells. Microscopy was performed at room temperature (21 °C) on a Nikon Eclipse Ti2 equipped with 488 nm and 561 nm lasers and dual ORCA-Fusion BT sCMOS cameras in the ALFM Facility (CSSB/DESY, Hamburg) using NIS-Elements v5.42.04. A water-immersion 60× objective (NA 1.27) was used. For each field of view, a 3-min movie was acquired with 100 ms exposure per channel at 2 s intervals (0.5 fps per channel) over 8-11 z-planes (Voxel size: 0.1083x0.1083x0.7 µm). To compensate for bleaching and/or change of fluorescent signal over time, we merged the two channels, as the sum of the signal from both channels. Images were cropped to remove uneven illumination. The cell segmentation and tracking were then performed in this created merged channel. The cell segmentation was performed with Cellpose(55) to provide cell identity to the tracking result. Tracking was done with TrackMate(56) using the Simple LAP tracker with linking max distance and gap-closing max distance of 1.0 µm and gap-closing max frame of 0. Only tracks from cells expressing Ent5 were kept, as defined by per-cell SNR and intensity thresholds (median SNR₂ ≥ 0.55, fraction of SNR₂ ≥ 0.70 ≥ 0.10, median f₂ ≥ 0.40) applied. Tracks touching acquisition boundaries were removed as timings cannot be determined for those. Each track was background-subtracted using fixed channel-specific values (Gga2: 100, Ent5: 100), then min–max normalized within track per channel. Track durations were binned into six intervals (0–10, 10–15, 16–20, 21–25, 26–30, >30 s), and a χ² test of independence was performed on the resulting contingency table (χ² = 54.43, df = 5, p = 1.71 × 10⁻¹⁰). For intensity overlays (Fig. 7d–e), an additional dynamic range filter was applied to exclude low-amplitude traces (defined as normalized range < 0.08) prior to alignment. Gga2 traces were aligned to their own intensity peak (t = 0), while Ent5 traces were aligned to the Gga2 peak from the corresponding track. Final overlays represent LOESS-smoothed dataset means (span = 0.2), with shaded bands indicating 95% confidence intervals estimated by per-timepoint bootstrapping. Sample sizes post-filtering: WT, n = 639; ΔHelix, n = 525.

## Data Availability

Ent5 x-ray structure was deposited to the PDB data bank with the 9S6D code. SAXS data was uploaded to SASDB with code SASDYA2. Microscopy images were uploaded to BioImage Archive with code S-BIAD2383. CD Spectra were uploaded to Zenodo with code (10.5281/zenodo.17523881).

## Code Availability

For the *in vivo* Gga2–Ent5 recruitment dynamics experiments, the fluorescence microscopy analysis pipeline is available at: https://git.embl.org/grp-cba/yeast-colocalization-tracking

## Acknowledgements

We acknowledge the staff of the EMBL P12/P14 (Aleksi Sutinen and Clement Blanchet for P12 and Michael Agthe for P14) beamlines as well as the Sample Preparation and Characterisation (SPC) facility of EMBL Hamburg at PETRA III (DESY, Hamburg) for assistance (Angelica Struve). We thank the Advanced Light and Fluorescent Microscopy (ALFM) facility at the Centre for Structural Systems Biology (CSSB) for support with light microscopy image recording and analysis. LAD was partly funded by EMBL Interdisciplinary Postdoc Programme (EIPOD3) under Marie Curie Actions COFUND 664726 and by the European Regional Development Fund (ERDF) through the Interreg Öresund-Kattegat-Skagerrak programme, project ‘Hanseatic League of Science (HALOS). GDB was funded by the EMBL International Ph.D. programme. CT has been supported by grant number 2020-225265 from the Chan Zuckerberg Initiative DAF, an advised fund of Silicon Valley Community Foundation.

## Author Contributions

G.D.B., J.S. and L.A.D. expressed and purified recombinant proteins and performed site-directed mutagenesis. G.D.B. produced and imaged GUVs with input from L.A.D. Mass photometry experiments were performed by J.S. Fluorescence microscopy and FRAP experiments were carried out by G.D.B. and J.S., with input from L.A.D. Circular dichroism spectroscopy was performed by J.S. L.A.D. performed stopped-flow light scattering experiments, and together with S.N. carried out stopped-flow time-resolved SAXS experiments. D.R.C. performed protein crystallization, X-ray diffraction data collection, and data processing. L.A.D. reconstructed phylogenetic trees and carried out AlphaFold3 modelling. Molecular biology of pRS315-Ent5-mScarlet, subsequent transformations and endogenous tagging of Gga2 with mNeonGreen were performed by K.V. *In vivo* Gga2–Ent5 recruitment dynamics were performed by L.A.D., with help from R.T. for data collection, and Z.H. and C.T. for data analysis. M.G.A. conceived and supervised the project together with L.A.D. L.A.D. and M.G.A. wrote the manuscript with input from all authors.

## Large language models

ChatGPT (https://chat.openai.com/) was used as an aid to correct written text. The authors take full responsibility for the manuscript content and code.

## Notes

### Competing Interest Statement

The authors have declared no competing interest.

