## Supplementary Information for "Clathrin adaptors drive phase separation in endocytosis and trafficking"

† - equally contributed

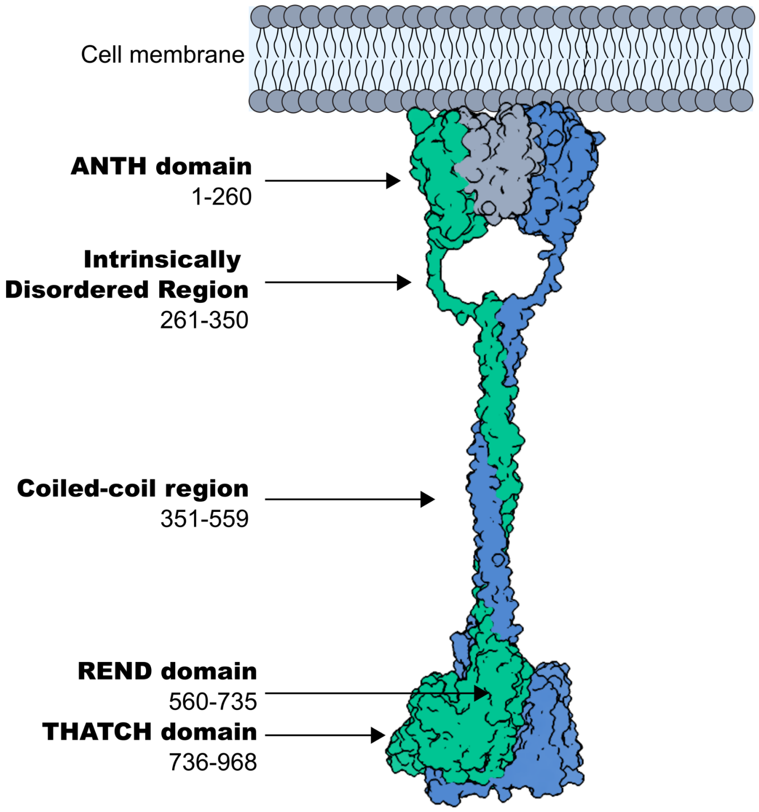


**Supplementary Figure 1: Sla2 structural representation scheme** (adapted from Draper-Barr et al., 2025). Sla2 dimerizes mainly through its coiled-coil region. In this representation, Sla2 monomers are shown in green and blue. Each Sla2 monomer contains an N-terminal ANTH membrane-binding domain (residues 1–260), which binds, in the presence of phosphatidylinositol 4,5-bisphosphate ($PIP_2$), to the ENTH domain of Ent1 (shown in grey). Two molecules of ANTH and two molecules of ENTH domains form a heterotetrameric membrane complex (the AENTH complex) (Lizarrondo et al., 2023). The ANTH domains of Sla2 are followed by an intrinsically disordered region (IDR; residues 261–350), a coiled-coil region (residues 351–559), a force-sensing domain (REND; residues 560–735), and an actin-binding domain (THATCH; residues 736–968).

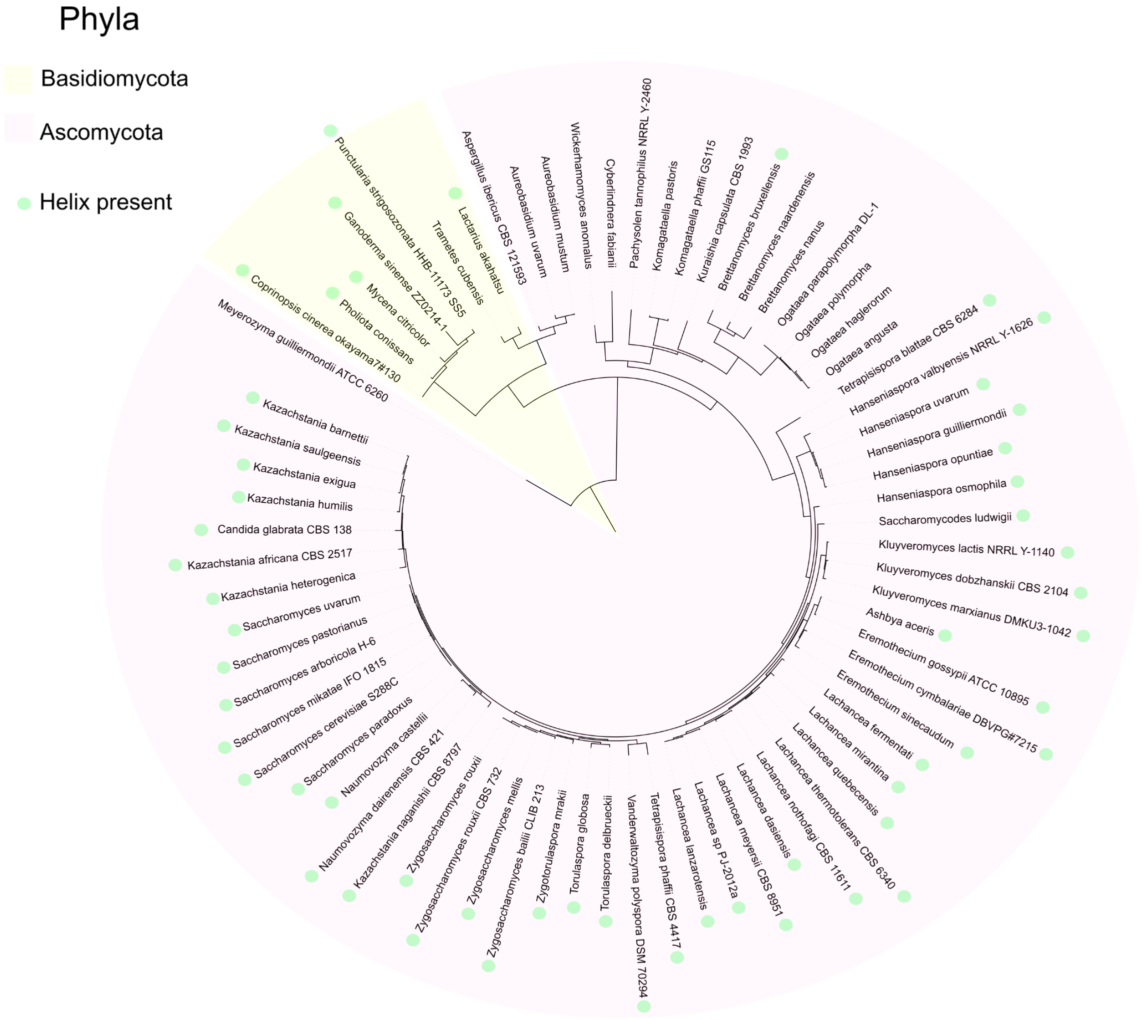


**Supplementary Figure 2. Distribution of Ent5 proteins and associated helix across Dykaria**. Green dots indicate species in which both Ent5 and the characteristic helix are detected by AF3.

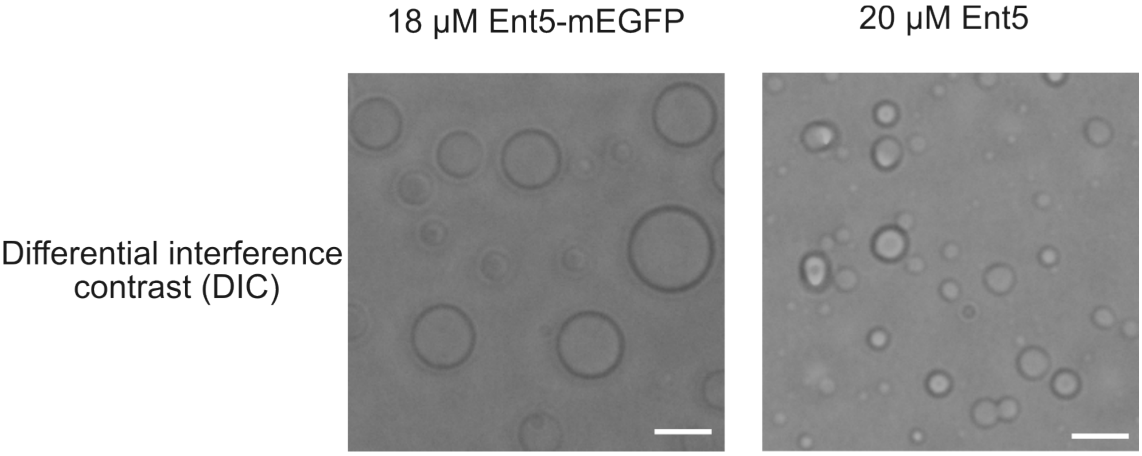


**Supplementary Figure 3. DIC Imaging of Ent5-mEGFP and Unlabeled Ent5.** DIC images of Ent5-mEGFP and unlabeled Ent5 demonstrate condensate formation in both samples, confirming that phase separation occurs independent of the mEGFP tag. White scale bars represent 5 µm. Relative to Figure 2.

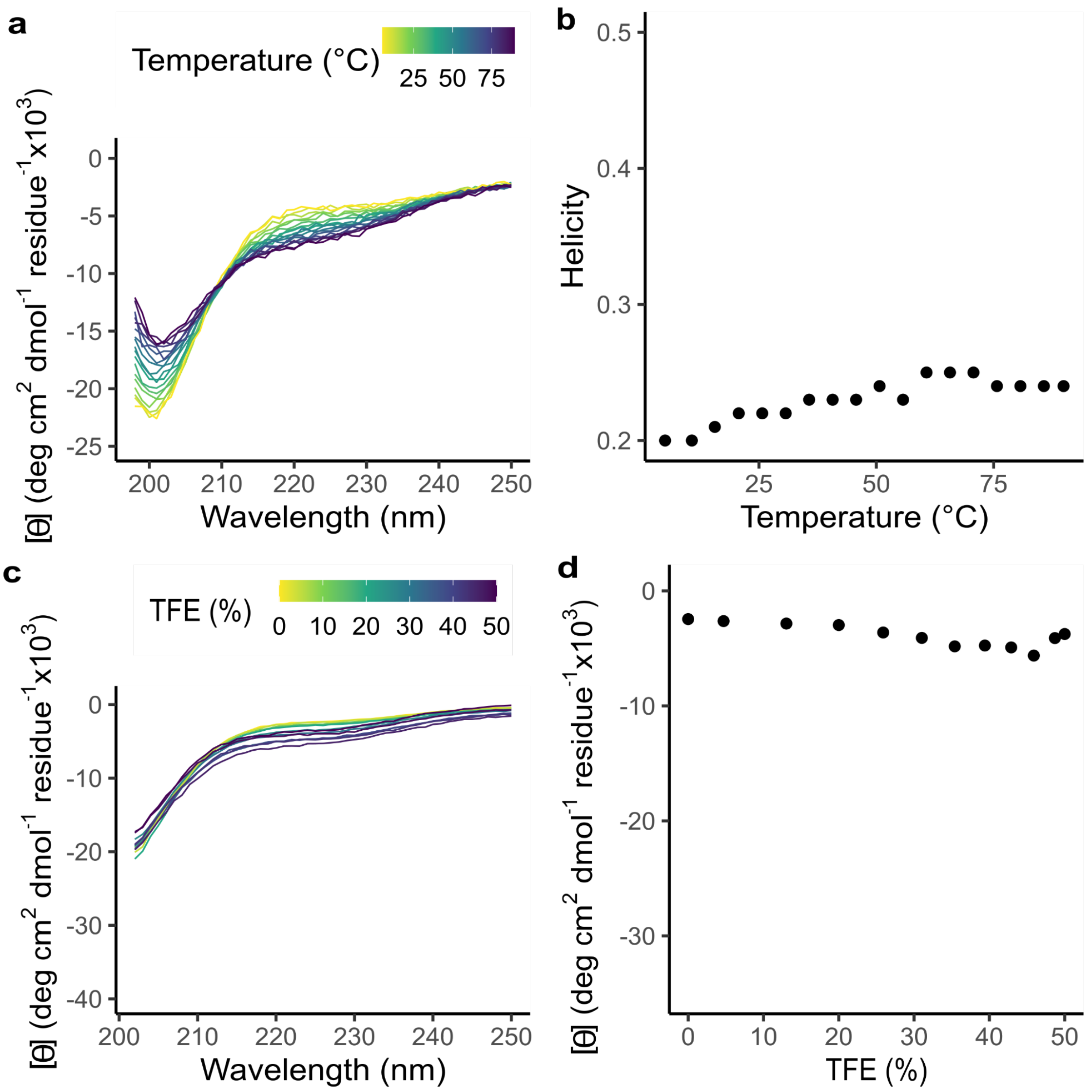


**Supplementary Figure 4. Circular dichroism of Ent5 6P Peptide.** a) Circular Dichroism (CD) spectra of the Ent5 Helix peptide (helix) were recorded over a temperature ramp from 5–90 °C. b)**.** The percentage contribution of helical secondary structure—calculated from the molar ellipticity spectra using the Peptide Helicity module of ChiraKit (1) —is plotted against temperature. c) Far UV CD spectra for the helix were recorded in the presence of TFE titrated from 0 to 50% (v/v) in 10 mM Tris (pH 7.5). d) Molar ellipticity at 222 nm plotted against TFE percentage.

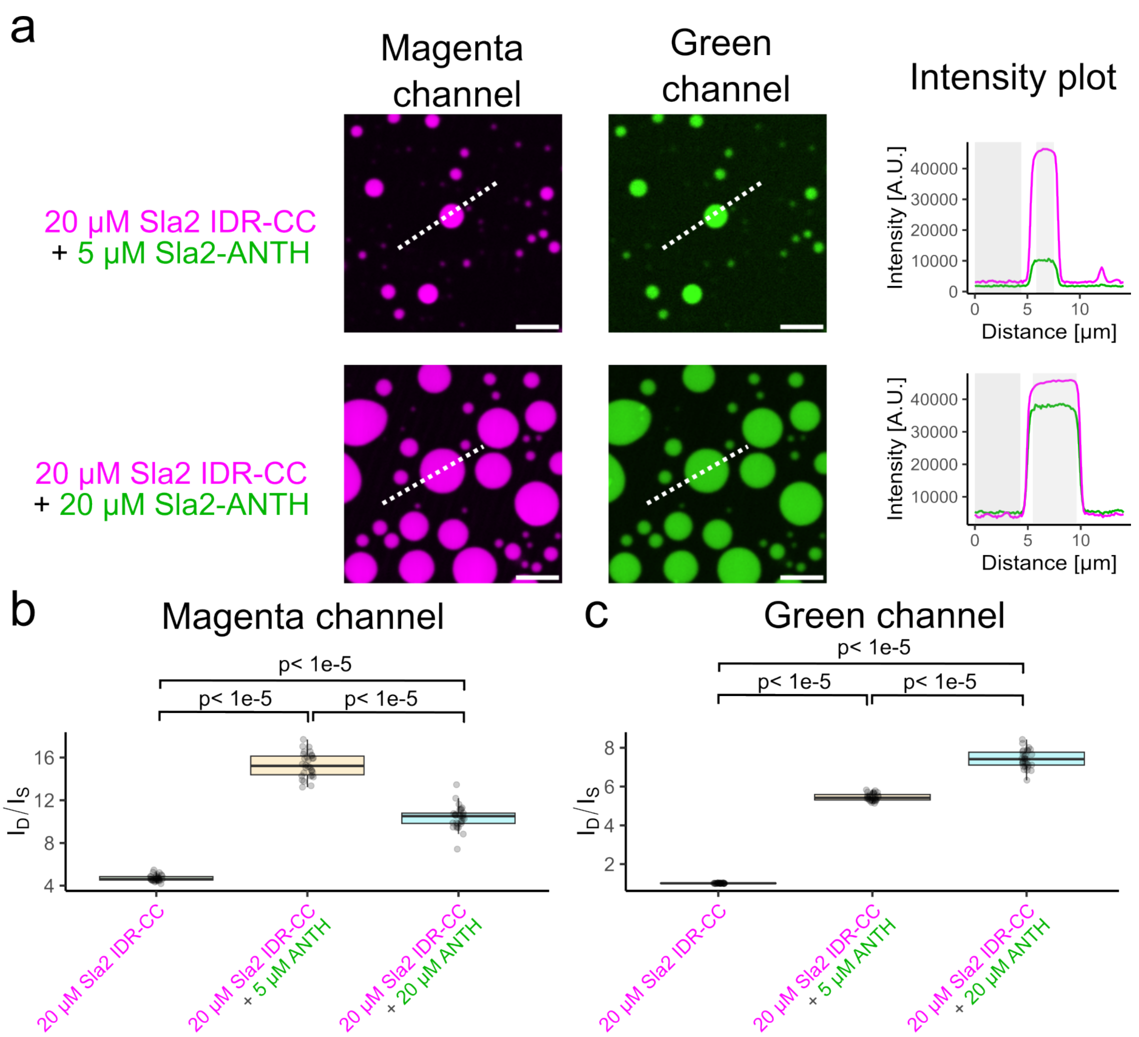


**Supplementary Figure 5. Representative Imaging and Intensity Quantification for Sla2 IDR-CC and Sla2 ANTH Interactions.**

a) Representative condensate images acquired under the same conditions as in Figure 5. Samples contained 20 µM Sla2 IDR-CC–mScarlet combined with either 5 µM or 20 µM Sla2–ANTH–mEGFP. In each image, a central line is drawn to quantify the difference in protein concentration between the droplet ($I_D$, intensity in the droplet) and the surrounding solvent ($I_S$, intensity in the solvent). White scale bars represent 5 µm.

b) Average intensity ratios ($I_D/I_S$) for the magenta channel (Sla2 IDR-CC–mScarlet) are plotted for the two conditions in panel a along with data for Sla2 IDR-CC alone (as reported in Figure 5). c) Average intensity ratios ($I_D/I_S$) for the green channel (Sla2–ANTH–mEGFP) are plotted for the two conditions in panel a along with data for Sla2 IDR-CC alone (from Figure 5). Statistical analyses were performed as described for Figure 5.

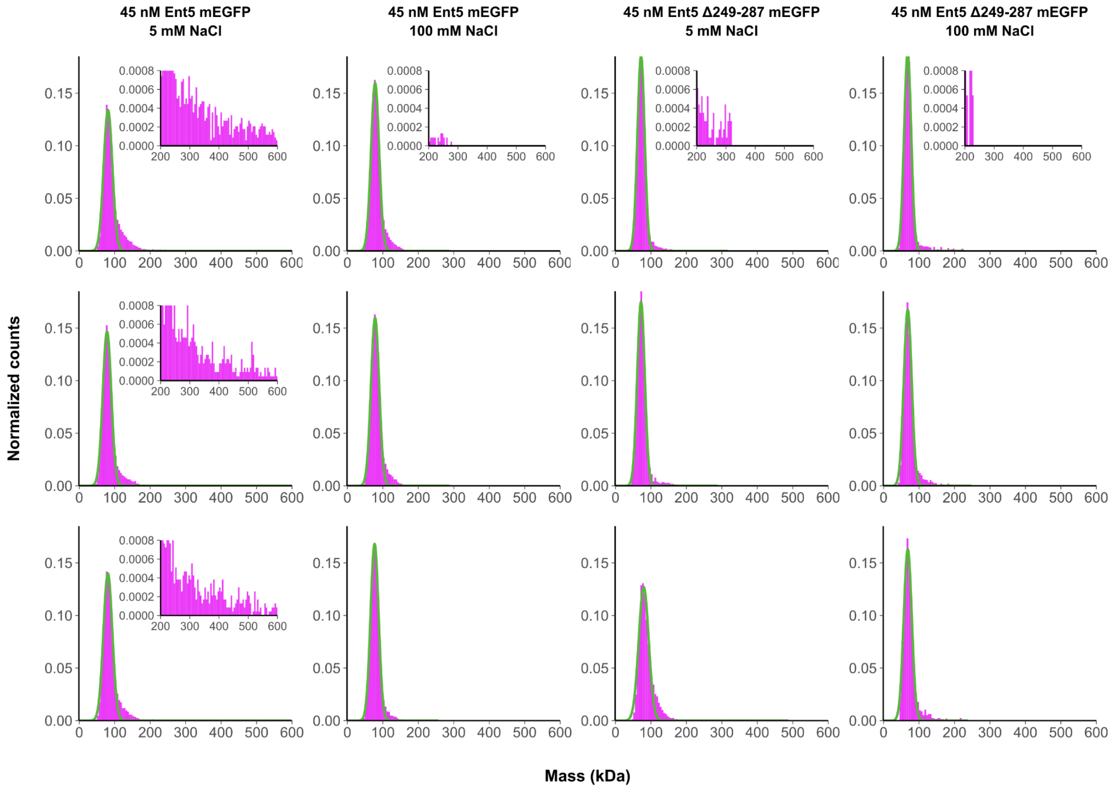


**Supplementary Figure 6. Characterization of Ent5WT-mEGFP Nanoclusters in Mass Photometry (MP).** Apparent molecular weight ($MW_{app}$) histograms obtained from MP measurements of 45 nM Ent5WT-mEGFP in 100 mM Tris (pH 7.5) with 100 mM NaCl and 5 mM NaCl, and of 45 nM Ent5Δ249-287-mEGFP in 100 mM Tris (pH 7.5) with 5 mM NaCl, from n = 3 independent experiments, histograms are shown in magenta while fitted mass are shown in green. The expected mass is 74 kDa for WT and 69 kDa for the Ent5Δ249-287-mEGFP. The inset magnifies the $MW_{app}$ range between 200 and 600 kDa at room temperature (RT).

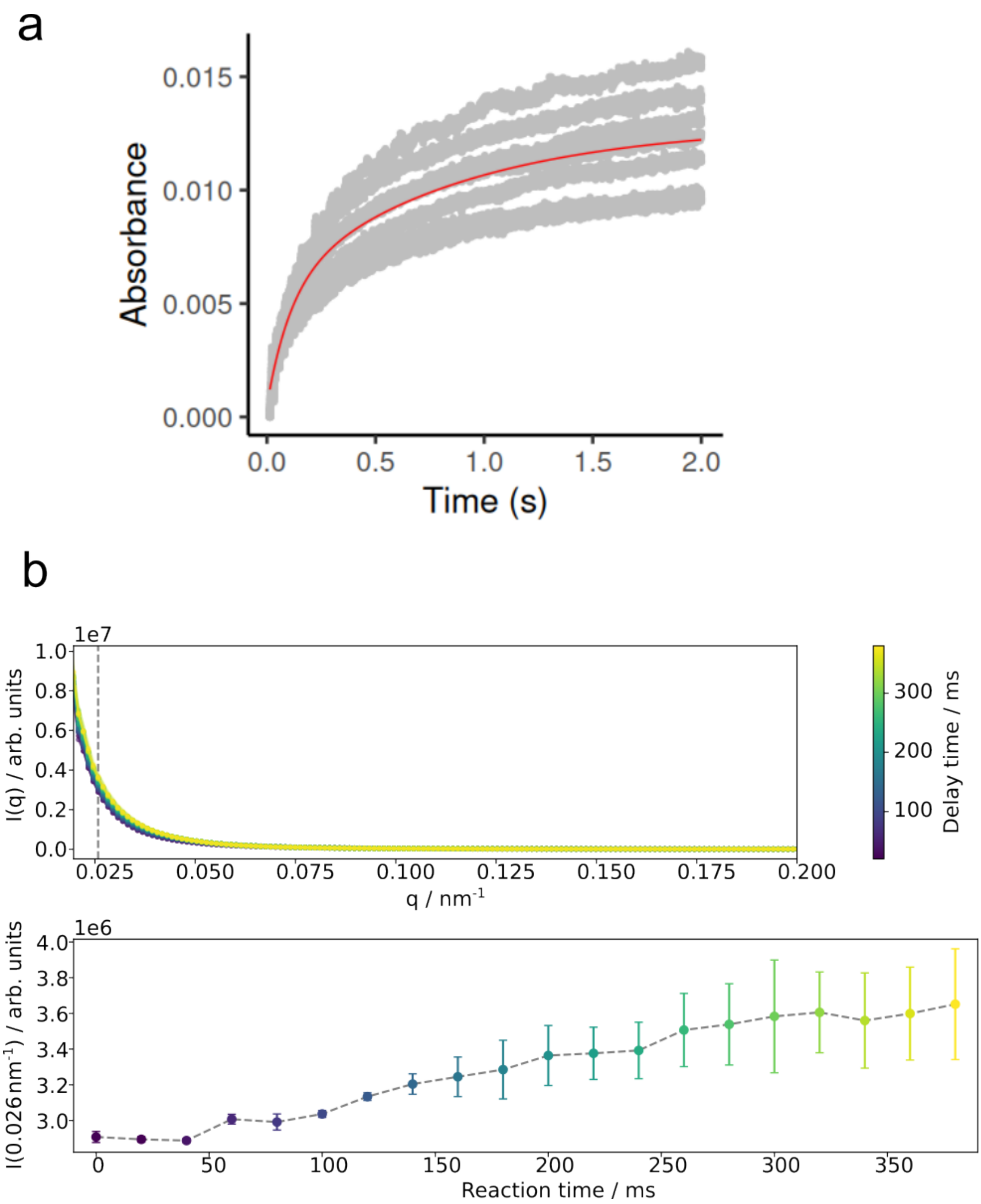


**Supplementary Figure 7. Light Scattering and SAXS.** a) Stopped-Flow Light Scattering. The red curve shows the fitting to a double exponential curve with time constants of $\tau_1$ (108±7) ms and $\tau_2$ (843±42) ms. b) Time-Resolved SAXS curves for different delay times (top) and a vertical slice at 0.026 $nm^{-1}$ (bottom).

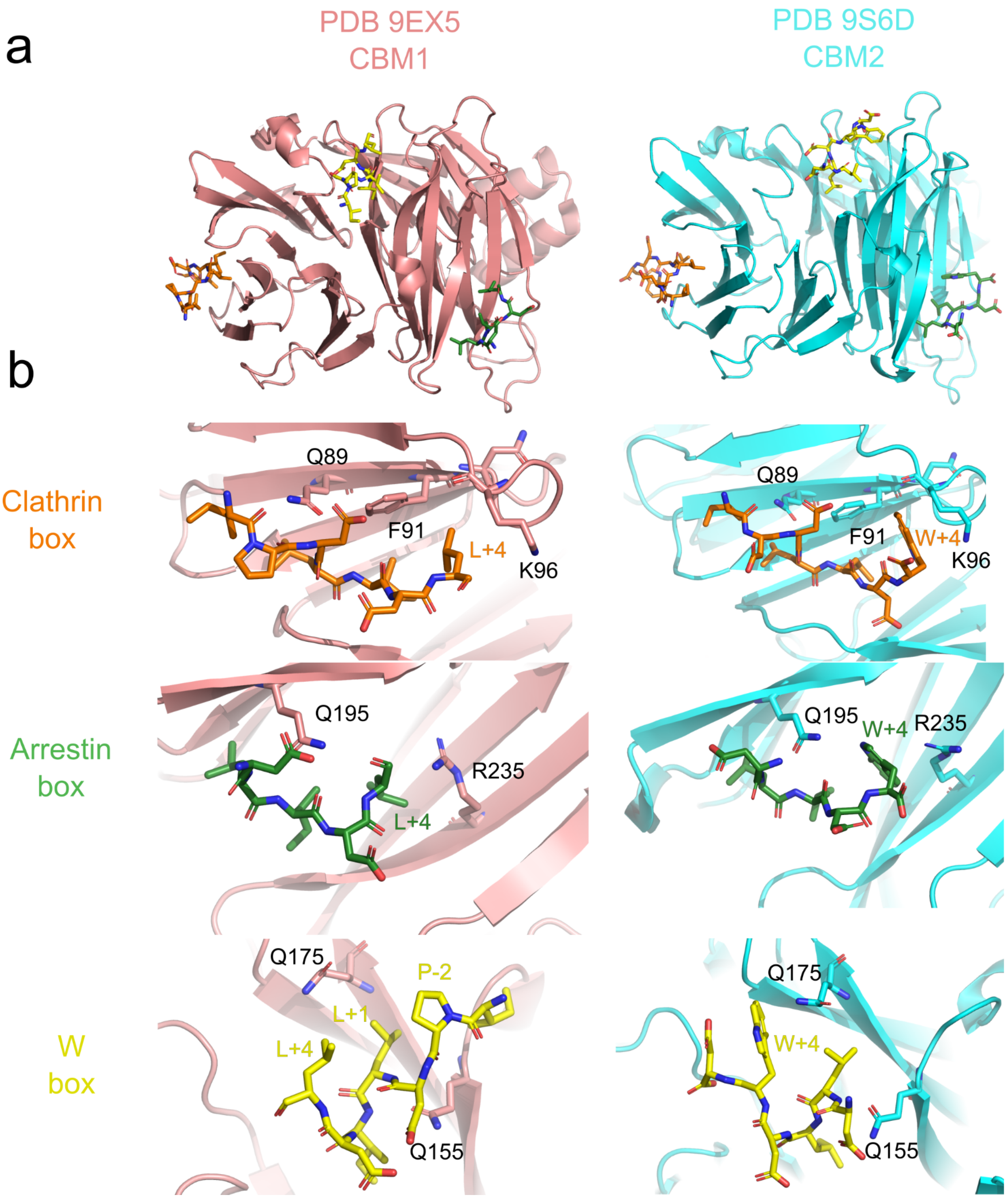


**Supplementary Figure 8. Crystal structure of NTD-CHC in complex with Ent5.2 peptide (LLDW).** a) Crystal structure of NTD-CHC bound to CBM1 of Ent5 (Left, PDB 5EX5). Crystal structure of NTD-CHC bound to CBM2 of Ent5 (Right, 9S6D). b) Close-up view of each binding box.

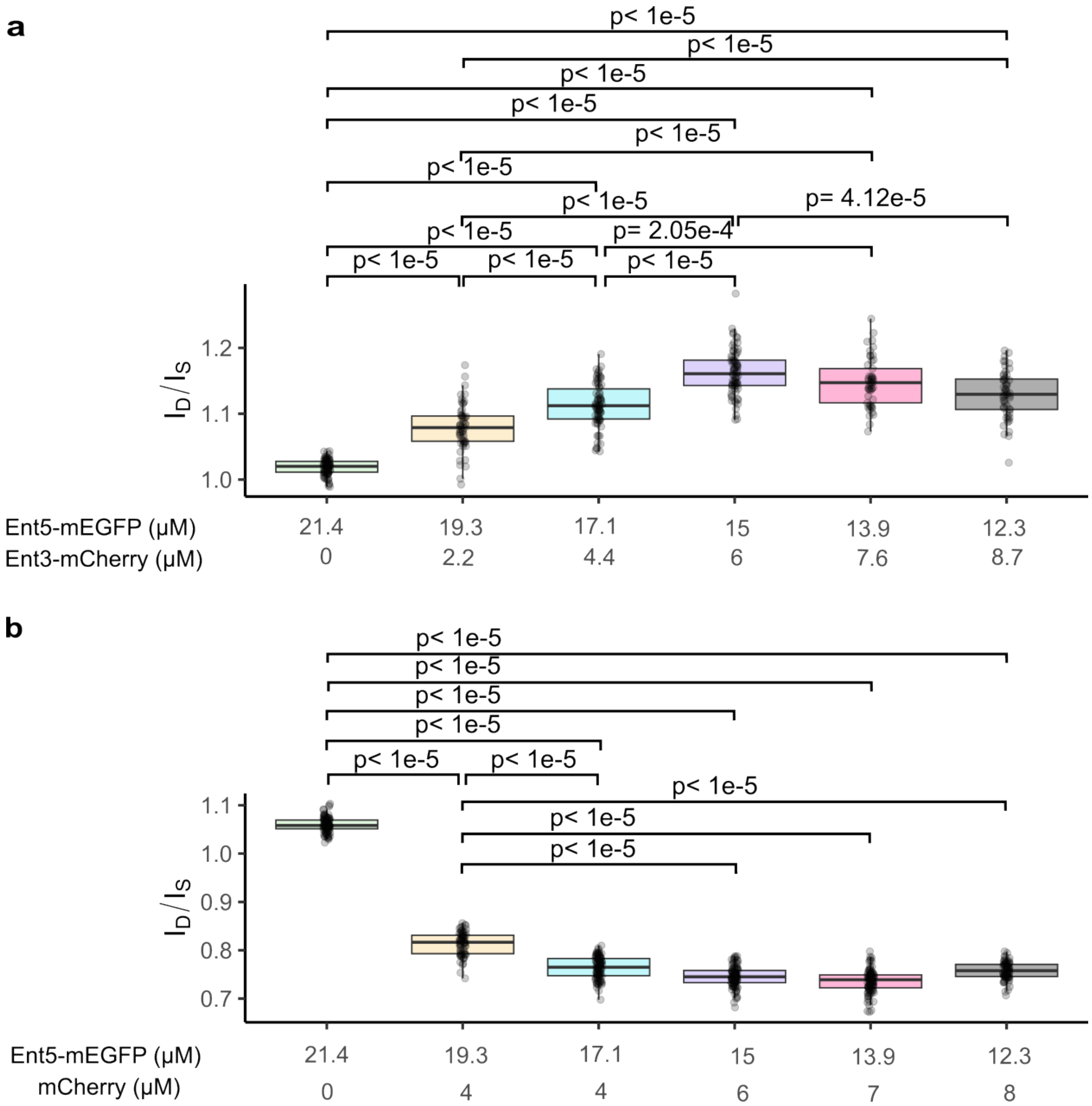


**Supplementary Figure 9. Quantitative Analysis of Ent3 Partitioning into Ent5 Droplets.** a) Partitioning coefficient of Ent3WT–mCherry into Ent5WT–mEGFP droplets is measured over a concentration range of 0–8.7 μM (Ent3WT–mCherry) and 12.3–20.4 μM (Ent5WT–mEGFP); n = 50 droplets were analyzed. b) Partitioning coefficient of mCherry into Ent5WT–mEGFP droplets is measured over a concentration range of 0–8.0 μM (mCherry) and 12.8–21.4 μM (Ent5WT–mEGFP); n = 70 droplets were analyzed. Statistical analyses were performed as described for Fig 5.

**Supplementary Table 1. IDR Regions and HotSpot residues for LLPS.**

| Name | IDR region | pLLPS | Droplet-promoting regions |
|---|---|---|---|
| Apl2 | 637-726 | 0.21 | 694-721 |
| Apl5 | 648-931 | 0.36 | 721-771;783-875;897-932 |
| CHC | - | 0.15 | 1623-1653 |
| CLC | 1-110 | 0.97 | 1-28;33-125 |
| Ede1 | 228-274, 367-597,884-1340 | 0.99 | 98-139;203-214;235-250;360-565;571-585;701-713;803-815;879-1029;1032-1054;1058-1147;1152-1202;1255-1286;1295-1340 |
| Ent1 | 306-454 | 0.97 | 108-118; 136-197; 241-226; 253-265; 290-369;425-452 |
| Ent2 | 416-613 | 1.00 | 128-212;227-257;286-415;423-613 |
| Ent3 | 173-408 | 0.99 | 158-272; 285-321; 340-378 |
| Ent5 | 206-411 | 0.97 | 199-411 |
| Gga1 | 1-18; 317-440 | 0.31 | 1-18;371-381;541-557 |
| Gga2 | 1-24; 335-470 | 0.73 | 256-269;335-351;354-465 |
| Sla1 | 130-253;412-497;619-656;718-1244 | 1.00 | 123-254; 397-510; 548-657; 718-858; 1027-1056; 1091-1111; 1129-1140; 1199-1228 |
| Sla2 | 260-351 | 0.57 | 278-370; 385-428; 593-604 |
| Swa2 | 1-140;250-367;516-551 | 0.84 | 1-98;295-363;451-464;513-560 |
| Yap1801 | 324-637 | 0.97 | 282-299;333-387;400-414;424-459;470-496;520-542;559-624 |
| Yap1802 | 325-568 | 0.89 | 255-438;442-454;556-568 |

**Supplementary Table 2. Buffer Composition Table**

| Buffer Name | Composition |
|---|---|
| PCR buffer | 40 mM Tris–HCl, pH 8.8 (25°C);<br>4 mM $MgCl_2$;<br>120 mM KCl;<br>20 mM $(NH_4)_2SO_4$;<br>0.02 mM EDTA;<br>0.2% Triton X-100;<br>8% glycerol;<br>0.005% Xylene Cyanol FF;<br>0.05% Orange G;<br>0.4 mM dNTPs;<br>0.04 U/µL Phu-Sso7d polymerase |
| HIC Buffer A (Ent5, Ent3) | 10 mM Tris–HCl pH 7.5<br>100 mM NaCl<br>0.25 mM TCEP<br>0.7 M $(NH_4)_2SO_4$<br>0.5 mM EDTA |
| HIC Buffer B (Ent5, Ent3) | 20 mM Tris–HCl pH 7.5<br>200 mM NaCl<br>0.5 mM TCEP<br>1 mM EDTA |
| HisTrap Buffer A1 (Ent5, Ent3, NTD-CHC) | 20 mM NaP pH 7.5<br>500 mM NaCl<br>12.5 mM Imidazole |
| HisTrap Buffer B1 (Ent5, Ent3, NTD-CHC) | 20 mM NaP pH 7.5<br>500 mM NaCl<br>250 mM Imidazole |
| Buffer A2 (Sla2, Ede1, CHC:1172-1574, CLC, Ent1-ENTH) | 30 mM HEPES pH 8<br>300 mM NaCl<br>5 % w/v glycerol |
| Buffer B2 (Sla2, Ede1, CHC:1172-1574, CLC, Ent1-ENTH) | 30 mM HEPES pH 8<br>300 mM NaCl<br>5 % w/v glycerol<br>200 mM Imidazole |
| Lysis Buffer (Ent5, Ent3) | 20 mM Sodium Phosphate NaP pH 7<br>200 mM NaCl<br>0.05% Tween-20, 2 mM MgCl2<br>0.5 mM TCEP<br>400 U DNAse I<br>Complete EDTA-Free protease inhibitor |

| | |
|---|---|
| | cocktail (Roche) |
| Lysis Buffer (Sla2, Ede1, CLC, $CHC^{1172-1574}$, Ent1-ENTH) | 30 mM HEPES pH 8<br>300 mM NaCl<br>5 % w/v glycerol<br>400 U DNAse I<br>Complete EDTA-Free protease inhibitor cocktail (Roche) |
| SEC Buffer 1 (Ent5, Ent3) | 100 mM Tris–HCl pH 7.5<br>150 mM NaCl<br>0.5 mM TCEP<br>1 mM EDTA |
| SEC Buffer 2 (CHC-NTD) | 50mM Tris–HCl pH 9<br>150 mM NaCl<br>0.5 mM TCEP |
| SEC Buffer 3 (Sla2, Ede1, Ent1-ENTH, CLC, $CHC^{1172-1574}$) | 30 mM HEPES pH 8<br>150 mM NaCl<br>0.5 mM TCEP |
| Dialysis Buffer 1 (Ent5, Ent3) | 20 mM TRIS pH 7.5<br>200 mM NaCl<br>0.5 mM TCEP<br>optional: 1 mM EDTA |
| Dialysis Buffer 2 (Sla2, Ede1, $CHC^{1172-1574}$, CLC, Ent1-ENTH) | 30 mM HEPES pH 8<br>150 mM NaCl<br>5 % w/v glycerol<br>1 mM DTT |

**Supplementary Table 3.** Primers used for QuikChange mutagenesis in both pETM-30-GST-NTD-CHC and pETM-11-Sumo3-Ent5-mEGFP

| Name | Protein | Mutation | Type | Sequence |
|---|---|---|---|---|
| K63E_Fw | NTD-CHC | K63E | Fw | GGCAATGAAGTGACAAGGGAGAATATGGGCGGTGATTCTGCTATCATG |
| K63E_Rev | NTD-CHC | K63E | Rev | CATGATAGCAGAATCACCGCCCATATTCTCCCTTGTCACTTCATTGCC |
| I87D+Q89A+K98E | NTD-CHC | I87D+Q89A+K98E | Fw | CAAATGGTACTgacGTGgccATATTTAATTTGGAAACTAAGAGCgagTTAAAG |
| I87D+Q89A+K98E | NTD-CHC | I87D+Q89A+K98E | Rev | CTTTAACTCGCTCTTAGTTTCCAAATTAAATATGGCCACGTCAGTACCATTTG |
| Q195A+I197T_Fw | NTD-CHC | Q195A+I197T | Fw | CTCAAAACAACGTAACATCTCCgccGCTaccGACGGTCATGTTG |
| Q195A+II197T_Rev | NTD-CHC | Q195A+I197T | Rev | CAACATGACCGTCGGTAGCGGCGGAGATGTTACGTTGTTTTGAG |
| K251E_Fw | NTD-CHC | K251E | Fw | GCTTCATTGCCTTCTCAATATCAAGAGGAAACTACCGATATTTTCTTTCC |
| K251E_Rev | NTD-CHC | K251E | Rev | GGAAAGAAAATATCGGTAGTTTCCTCTTGATATTGAGAAGGCAATGAAGC |
| F26A_Fw | NTD-CHC | F26A | Fw | GGAATTTCCCCTCAATTCCTTGACGCCAGATCAACTACTTTCGAG |
| F26A_Rev | NTD-CHC | F26A | Rev | CTCGAAAGTAGTTGATCTGGCGTCAAGGAATTGAGGGGAAATTCC |

| | | | | |
|---|---|---|---|---|
| Q155A_Fw | NTD-CHC | Q155A | Fw | CCTTGAGACACGCTAACTTGAACAATACCgccATTATCAATTTTGTGGCTAACAAAAACC |
| Q155A_Rev | NTD-CHC | Q155A | Rev | GGTTTTTGTTAGCCACAAAATTGATAATGGCGGTATTGTTCAAGTTAGCGTGTCTCAAGG |
| Delta249-287_Fw | Ent5 | Delta Helix | Fw | GTGACCAACTTCAATGTTCCTGTGGAAACAGAAGATAGCATCCCTGACTT |
| Delta249-287_Rev | Ent5 | Delta Helix | Rev | AAGTCAGGGATGCTATCTTCTGTTTCCACAGGAACATTGAAGTTGGTCAC |
| 211-411_Fw | Ent5 | IDR + Helix | Fw | caccattgacgtgttccagcaacagaccggtACGTCTGCAAGTTACGATAACATCG |
| 211-411_Rev | Ent5 | IDR + Helix | Rev | CGATGTTATCGTAACTTGCAGACGTaccggtctgttgctggaacacgtcaatggtg |
| 1-210_Fw | Ent5 | ENTH | Fw | CCATTTATAATGCCAACCAGATTTCAggatccgtgagcaagggcgaggagctgttc |
| 1-210_Rev | Ent5 | ENTH | Rev | gaacagctcctcgcccttgctcacggatccTGAAATCTGGTTGGCATTATAAATGG |

**Supplementary Table 4.** X-ray data collection and refinement statistics. Statistics for the highest-resolution shell are shown in parentheses

| | |
|---|---|
| PDB code | 9S6D |
| Wavelength (Å) | 0.9763 |
| Resolution range (Å) | 35.56 - 1.7 (1.761 - 1.7) |
| Space group | C 1 2 1 |
| Unit cell (Å,°) | 96.341 129.881 74.532 90.0 96.673 90.0 |
| Total reflections | 685795 (69298) |
| Unique reflections | 97208 (9575) |
| Multiplicity | 7.1 (7.2) |
| Completeness (%) | 97.47 (96.15) |
| Mean I/sigma(I) | 12.88 (1.69) |
| Wilson B-factor | 32.58 |
| R-merge | 0.07639 (1.21) |
| R-meas | 0.08265 (1.303) |
| R-pim | 0.03115 (0.4801) |
| CC1/2 | 0.997 (0.716) |
| CC* | 0.999 (0.913) |
| Reflections used in refinement | 97197 (9569) |
| Reflections used for R-free | 4860 (479) |
| R-work | 0.1830 (0.3425) |
| R-free | 0.2114 (0.3606) |
| CC(work) | 0.965 (0.775) |

| | |
|---|---|
| CC(free) | 0.959 (0.757) |
| Number of non-hydrogen atoms | 6471 |
| macromolecules | 6080 |
| solvent | 391 |
| Protein residues | 777 |
| RMS(bonds) | 0.011 |
| RMS(angles) | 1.84 |
| Ramachandran favored (%) | 97.63 |
| Ramachandran allowed (%) | 2.37 |
| Rotamer outliers (%) | 1.19 |
| Clashscore | 2.31 |
| Average B-factor (Å²) | 42.44 |
| Macromolecules (**Å²)** | 42.42 |
| Solvent (Å²) | 42.69 |
